# Canonical Strigolactones Are Not the Tillering-Inhibitory Hormone but Rhizospheric Signals in Rice

**DOI:** 10.1101/2022.04.05.487102

**Authors:** Shinsaku Ito, Justine Braguy, Jian You Wang, Akiyoshi Yoda, Valentina Fiorilli, Ikuo Takahashi, Muhammad Jamil, Abrar Felemban, Sho Miyazaki, Teresa Mazzarella, Akihisa Shinozawa, Aparna Balakrishna, Lamis Berqdar, Chakravarty Rajan, Shawkat Ali, Imran Haider, Yasuyuki Sasaki, Shunsuke Yajima, Kohki Akiyama, Luisa Lanfranco, Matias Zurbriggen, Takahito Nomura, Tadao Asami, Salim Al-Babili

## Abstract

The plant hormones strigolactones (SLs) regulate shoot branching and mediate the communication with symbiotic mycorrhizal fungi, but also with noxious root parasitic weeds, such as *Striga* spp. SLs derive from carlactone (CL) and are divided structurally into canonical and non-canonical SLs. However, the questions about particular biological functions of the two groups and the identification of the SL that inhibits shoot branching are still unanswered, hampering targeted modification of SL pattern towards improving plant architecture and resistance against *Striga*. Here, we reported that 4-deoxyorobanchol (4DO) and orobanchol, the two canonical SLs in rice, do not have major role in determining rice shoot architecture. CRISPR/Cas9 mediated *Osmax1-900* mutants, lacking these two SLs, do not show the high tillering and dwarf phenotype typical for SL-deficient plants. However, the absence of 4DO and orobanchol in root exudates significantly decreased their capability in inducing *Striga* seed germination, while caused only a delay in root colonization by mycorrhizal fungi. To confirm the genetic evidence, we used the SL-biosynthesis inhibitor TIS108. Our results showed that TIS108 is a MAX1-specific inhibitor that lowers 4DO and orobanchol synthesis, conferring a resistance to *Striga* without a severe impact on rice architecture. Hence, our work uncovers the specific function of canonical SLs as rhizospheric signals and paves the way for establishing chemical and genetic based approaches for combating the root parasitic weeds, by targeted depletion of their release.

---

Strigolactones (SLs) are carotenoid-derived hormones characterized by an enol-ether bridge connecting a lactone ring (D-ring; Fig.S1) (Koichi Yoneyama et al. 2018) in *R* configuration to a structurally variable second moiety that consists of a tricyclic lactone ring (ABC-ring) in canonical SLs, while non-canonical SLs have variable structures based on a β-ionone ring (A-ring) (Fig.S1) (Al-Babili and Bouwmeester 2015). SLs are a major determinant of plant architecture; one of their key involvement amid several other biological processes. Among other phenotypes, mutants affected in SL biosynthesis are characterized by increased branching/tillering, shorter shoots (dwarf), and decreased primary root length (Morris et al. 2001; Gomez-Roldan et al. 2008; Al-Babili and Bouwmeester 2015).

In addition, when exposed to nutrients deficiency, particularly phosphate, plant roots release SLs to attract arbuscular mycorrhizal fungi (AMF). The latter establish the AM symbiosis, the most common type of plant mutualist association that significantly increases the uptake of nutrients and water from the soil (Akiyama, Matsuzaki, and Hayashi 2005; Marzec 2016; Lanfranco, Fiorilli, and Gutjahr 2018). However, canonical SLs were first discovered as the host-derived signals that stimulate seed germination in root parasitic weeds, such as *Orobanche* and *Striga spp*. (Cook et al. 1966). During their evolution, these obligate parasites have acquired the ability to utilize SLs as signal to coordinate their development with the presence of an available host in the close vicinity (Toh et al. 2015). Infestation by root parasitic plants, such as *Striga hermonthica*, is a severe problem for agriculture and a major threat for global food security, particularly in Africa, where it causes more than US$7 billion annual losses in cereal production (Mohamed et al. 2006; Parker 2012).

The availability of high-branching mutants of monocot and dicot plant species (Snowden et al. 2005; Stirnberg, Furner, and Ottoline Leyser 2007; Koltai et al. 2010; Arite et al. 2007; Cardoso et al. 2014) paved the way for discovering the hormonal function of SLs and enabled later the elucidation of their biosynthesis. SL biosynthesis starts with the reversible isomerization of all-*trans-* into 9-*cis*-β-carotene, catalyzed by DWARF27 (Abuauf et al. 2018; Alder et al. 2012; Bruno and Al-Babili 2016). It is followed by cleavage and rearrangement reactions, mediated by the CAROTENOID CLEAVAGE DIOXYGENASE 7 and 8 (CCD7/D17 and CCD8/D10), which yield carlactone (CL), the core intermediate of SL biosynthesis (Fig. S2) (Alder et al. 2012; Bruno and Al-Babili 2016). The discovery of CL unraveled the presence of the non-canonical SLs that were unknown before. Indeed, different modifications of CL, which are catalyzed by cytochrome P450 monooxygenases (CYP), in particular MORE AXILLARY GROWTH1 (MAX1) from the CYP711A clade, and other enzymes, give rise to the structural diversity of the more than 30 natural canonical and non-canonical SLs (Booker et al. 2005; Cardoso et al. 2014; Lazar and Goodman 2006; Wakabayashi et al. 2019).

Rice contains five *MAX1* homologs - *Os01g0700900* (*OsMAX1-900*), *Os01g0701400* (*OsMAX1-1400*), *Os01g0701500* (*OsMAX1-1500*), *Os02g0221900* (*OsMAX1-1900*) and *Os06g0565100* (*OsMAX1-5100*) (Nelson and Werck-Reichhart 2011; R. J. Challis et al. 2013) – with a truncated *OsMAX1-1500* in the Nipponbare cv. (Richard J. Challis et al. 2013). *In vitro* studies and transient expression in *Nicotiana benthamiana* showed that all functional Nipponbare OsMAX1 enzymes (OsMAX1-900, OsMAX1-1400, OsMAX-1900, and OsMAX1-5100) can convert CL into carlactonoic acid (CLA) that is transformed into the canonical SLs 4-deoxyorobanchol (4DO), and then orobanchol by sequential action of OsMAX1-900 and OsMAX1-1400 (Fig. S2) (Zhang et al. 2014; Kaori Yoneyama et al. 2018).

In this work, we investigated the biological function of canonical SLs in rice. For this purpose, we generated two bi-allelic homozygous *OsMAX1-900* knockout lines (*Os900*-KO: *Os900-32* and *-34*) disrupted in the biosynthesis of 4DO and orobanchol through introducing CRISPR/Cas9-induced deletion, point mutation and frameshift mutations (Fig. 1A). We first quantified 4DO and orobanchol in roots and root exudates of hydroponically grown and phosphate-starved mutants by Liquid Chromatography Tandem-Mass Spectrometry (LC-MS/MS) (Fig. 1B; Fig. S3A-B). 4DO and orobanchol were undetectable in both lines, confirming *in planta* the role of OsMAX1-900 as the rice 4DO synthase (Kaori Yoneyama et al. 2018) and that 4DO is the exclusive precursor of orobanchol in rice. Besides the absence of 4DO and orobanchol, exudates of the mutant lines showed a decrease of more than 96% in the level – and absent in rice root tissues - of a non-canonical SL tentatively identified as 4-oxo-MeCLA (4-oxo-methyl-carlactonoate) (Fig. S3C)), which was previously described as methoxyl-5-deoxystrigol isomer (Yoneyama et al., 2018). Based on the ion peak characteristic of the D-ring at 97.028, we also identified a novel SL, CL+30 with a molecular formula C_19_H_24_O_5_ (m/z 333.16989 as positive ion [M + H]^+^, calcd. for m/z 333.16965), which was present at high levels in the *Os900* mutants (Fig. S3C). Feeding *Os900*-34 seedlings with [^13^C]-labeled CL confirmed that CL+30 is a downstream product of CL (Fig. S4); however, the enzyme responsible for the production of this metabolite remains elusive, as we did not get any hint for the involvement of OsMAX1s from the transcript analysis (Fig. S5). The higher accumulation of CL+30 in *Os900*-KO lines (Fig. S3B-C) indicated that it might be a substrate of OsMAX1-900. We confirmed this assumption by expressing OsMAX1-900 in yeast cells and feeding them with a CL+30 containing fraction. After incubation and LC-MS/MS analysis, we detected a reduction in CL+30 content and its conversion into a novel metabolite eluting at 6.1 min (m/z 347 in positive-ion mode and 345 in negative-ion mode), corresponding to CL+30+14 Da (CL+30+14) (Fig. S6). As OsMAX1-900 catalyzes the carboxylation of CL, we expected the arising metabolite to contain a carboxyl group. Therefore, we methylated the novel OsMAX1-900 product by diazomethane, which gave rise to a derivative with m/z 361 in positive ion mode and fragment pattern and retention time (9.1 min), which are characteristic for the tentative 4-oxo-MeCLA (Fig. S7). Given that OsMAX1-900 catalyzes the oxidation at the C19 position, we assumed that CL+30 corresponds to 4-oxo-19-hydroxy-CL (Fig. S8).

**FIGURE 1.**
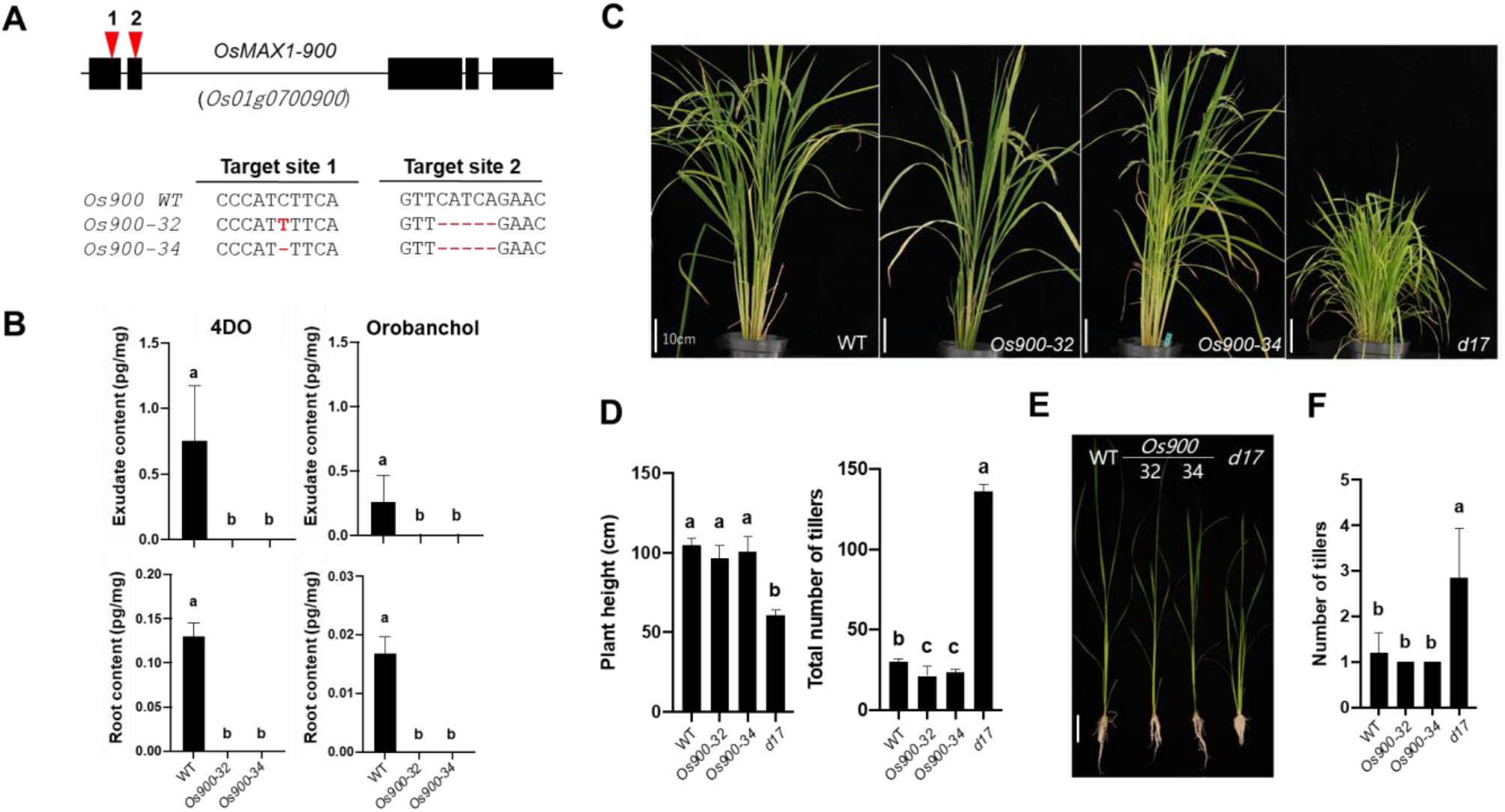
Generation of the *Os900*-knockout lines by CRISPR/Cas9 system (**A**) The structure of the *Os900* gene and the sequences of the two CRISPR/Cas9 target sites indicated by red arrows. Details of the CRISPR-mediated mutations of the two KO lines, *Os900-*32 and -34, are reported. (**B**) Analysis of canonical SLs, 4DO and orobanchol, in root exudates of *Os900*-KO lines grown under constant low Pi conditions. The data are presented as means ± SD from 5 samples. Means not sharing a letter in common differ significantly at *P*_0.05_. (**C** and **D**) Shoot phenotypes of WT, Os*900-*KO lines, and *d17* mutant grown in soil and hydroponic culture under +Pi conditions (E and F). The data are presented as mean ± SD for the number of biological replicates (C and D, 5≤n≤7 for WT, *Os900-32* and *-34*, n=3 for *d17*; E and F, 4≤n≤8).

Next, we phenotyped the growth and development of the *Os900*-KO lines, in comparison with WT and the high-tillering SL-deficient *d17* mutant (Butt et al. 2018). In soil and under normal growth conditions (+Pi), shoots of mature *Os900*-KO plants did not differ significantly from WT, in contrast to *d17* that showed the characteristic dwarfism and extreme high-tillering (Fig. 1C-E). Interestingly, *Os900*-KO lines had even less tillers, compared to WT (an average of 30 tillers for WT *vs* 21.4 and 23.6 tillers for *Os900-32* and *-34*, respectively) (Fig. 1D; Fig. S9A). *Os900*-KO mutants, grown in rhizotrons under normal conditions, showed a higher number of crown roots and root area, compared to WT (Fig. S9B-C). When hydroponically grown under different conditions (+Pi, -Pi, and low Pi), we did not detect common significant differences in shoot and root phenotype between the two mutants and the WT; with the exception of shorter shoots, lighter shoot, and root biomass under both +Pi and -Pi conditions (Fig. 1E; Fig. S10). Nevertheless, we did not detect pronounced morphological alterations, which are characteristic for SL deficient mutants (*d10* and *d17*), in the *Os900*-KO mutants in all three experiments, indicating that (1) canonical SLs are not major regulators of rice architecture and (2) the *Os900*-KO mutant lines still maintain a normal SL hormone homeostasis. To check the first assumption, we fed hydroponically grown *d17* seedlings with different concentrations of 4DO (0 nM, 1 nM, 10 nM, 100 nM, and 1000 nM) under normal conditions, using 1000 nM *rac-*GR24 (SL analog) as a positive control (Jamil et al., 2018) (Fig. S11A), and determined the effect of the treatment on their phenotype. We observed a decrease in tillering only at higher concentrations (100 and 1000 nM, Fig. S11B), which are much higher than endogenous SL levels (usually at picomole level under nutrient deficiency conditions). For the second hypothesis, we treated *d17* and *Os900*-KO mutants with 2.5 μM zaxinone, a growth-promoting apocarotenoid that requires intact SL biosynthesis and perception for its activity (Wang et al. 2019), with and without 1 μM *rac*-GR24 (Fig. S12). As expected, the application of zaxinone increased the *d17* root length only when combined with GR24. In contrast, zaxinone alone significantly enhanced the root and shoot length of *Os900*-KO lines (Fig. S13), suggesting that SL hormone biosynthesis and signaling are working properly in the absence of 4DO and orobanchol. In conclusion, these data suggested that canonical SLs are not responsible for regulating the tiller number in rice, which is in line with a recently published study on the role of orobanchol in tomato (Wakabayashi et al. 2019). Moreover, our results indicated that non-canonical SLs are the SL hormone regulating shoot architecture. This was further supported by the absence of canonical SLs and the presence of CL+30, as CL+30 was the only SL detected in the root-shoot junction (area where the tillers emerge) of *Os900* mutants, which do not have a shoot architecture-related phenotype (Fig. S14).

Next, we investigated the role of canonical SLs as rhizospheric signals. First, we estimated the colonization of *Os900*-KO roots by the AMF *Rizophagus irregularis* after 10-, 20- and 35-days post inoculation (dpi). For this purpose, we used the transcript abundance of *OsPT11*, a plant marker gene for a functional AM symbiosis (Guimil et al. 2005). At 10 dpi, there was a delay in colonization of *Os900*-KO roots compared to WT roots; whereas, at 20 and 35 dpi the colonization of *Os900* mutants was comparable to the WT (Fig. 2A). No other phenotypic differences were observed in intraradical fungal structures (Fig. 2B) and in plant traits (Fig. S15). Additionally, application of *Os900*-KO root exudates to *Gigaspora margarita* spores led to an induction of germination rate in analogy to *rac*-GR24 (Fig. S16), suggesting that exudates of *Osmax1-900* mutants still have non-canonical SLs at a sufficient level to sustain AMF germination in the absence of canonical SLs.

**FIGURE 2.**
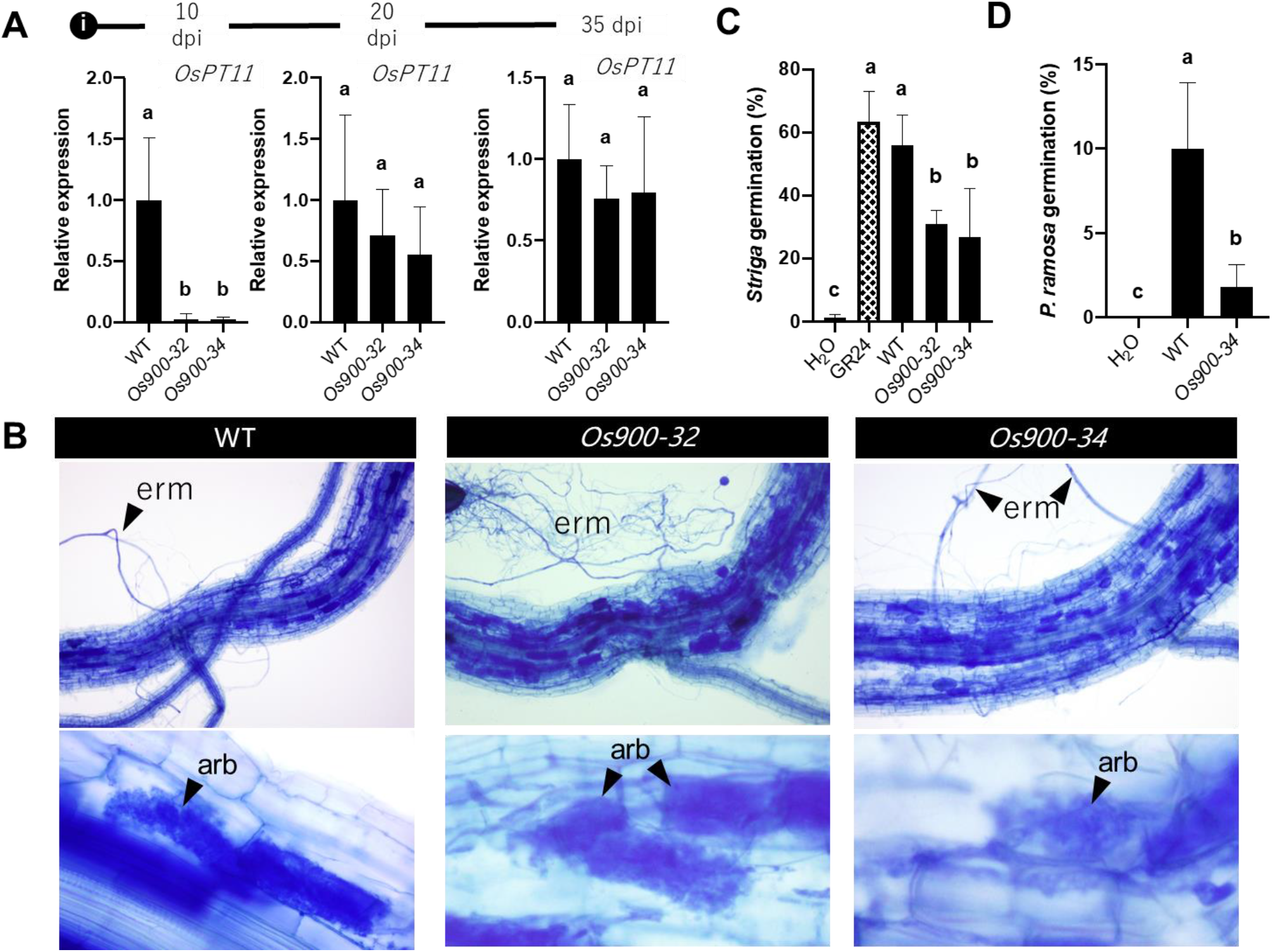
Effect of *Os900*-KO lines on the arbuscule formation (**A, B**) and the germination of root parasitic weeds (**C**, *Striga*, and **D**, *Phelipanche*). The values are represented as the mean ± SD for the number of biological replicates (A and B, n=4; C, 2<n<4; and D, n=3). The statistical significance is determined by one-way ANOVA and Tukey’s multiple comparison test. Arbuscule formation of *R. irregularis* was quantified by measuring the expression of marker gene (*OsPT11*) (A). (B) Arbuscule formation at 35 dpi.

We then tested the germination activity of *Os900* KO-lines root exudates on *Striga hermonthica* and *Phelipanche ramosa* seeds and observed more than 50% decrease in the germination of both parasitic species, compared to WT exudates (Fig. 2C-D; Fig. S17). This indicates that 4DO and orobanchol are important cues for parasitic seed germination, especially 4DO that was shown to be a stronger germination signal than orobanchol (Ueno et al. 2011). Hence, we can conclude that the two rice canonical SLs, 4DO and orobanchol, are rhizospheric signals important for the interaction with root parasitic plants and that decreasing their biosynthesis or even completely knocking it out is highly desired for reducing the damage caused by *Striga* and other root parasitic plants, without causing severe plant architectural changes. However, modulation of SL contents by genetic modifications requires years of development; while chemically-induced inhibition of their biosynthesis may lead much faster to rice plants lacking 4DO and orobanchol. Therefore, we set out to identify chemical(s) that inhibit canonical SL biosynthesis in rice.

TIS108 is an inhibitor of SL biosynthesis, which contains a 1*H*-1,2,4-triazole moiety (Fig. 3A) that can bind to the heme iron of P450s, such as MAX1 enzymes, and potentially impede their function(s) (Ito et al., 2011). Indeed, it inhibited the conversion of CL to CLA to 4DO by OsMAX1-900 (IC_50_ = 0.15 µM, for both conversions), and of 4DO to orobanchol by OsMAX1-1400 (IC_50_ = 0.02 µM) (Fig. 3A), when added to assays with microsomes prepared from yeast cells overexpressing the corresponding MAX1 enzyme. We could not determine whether TIS108 also affects the activity of OsMAX1-5100 and -1900, as we did not detect the sufficient conversion of CL to CLA, neither with native nor with codon-optimized OsMAX1-5100 and -1900 in yeast microsomes (Fig. S18).

**FIGURE 3.**
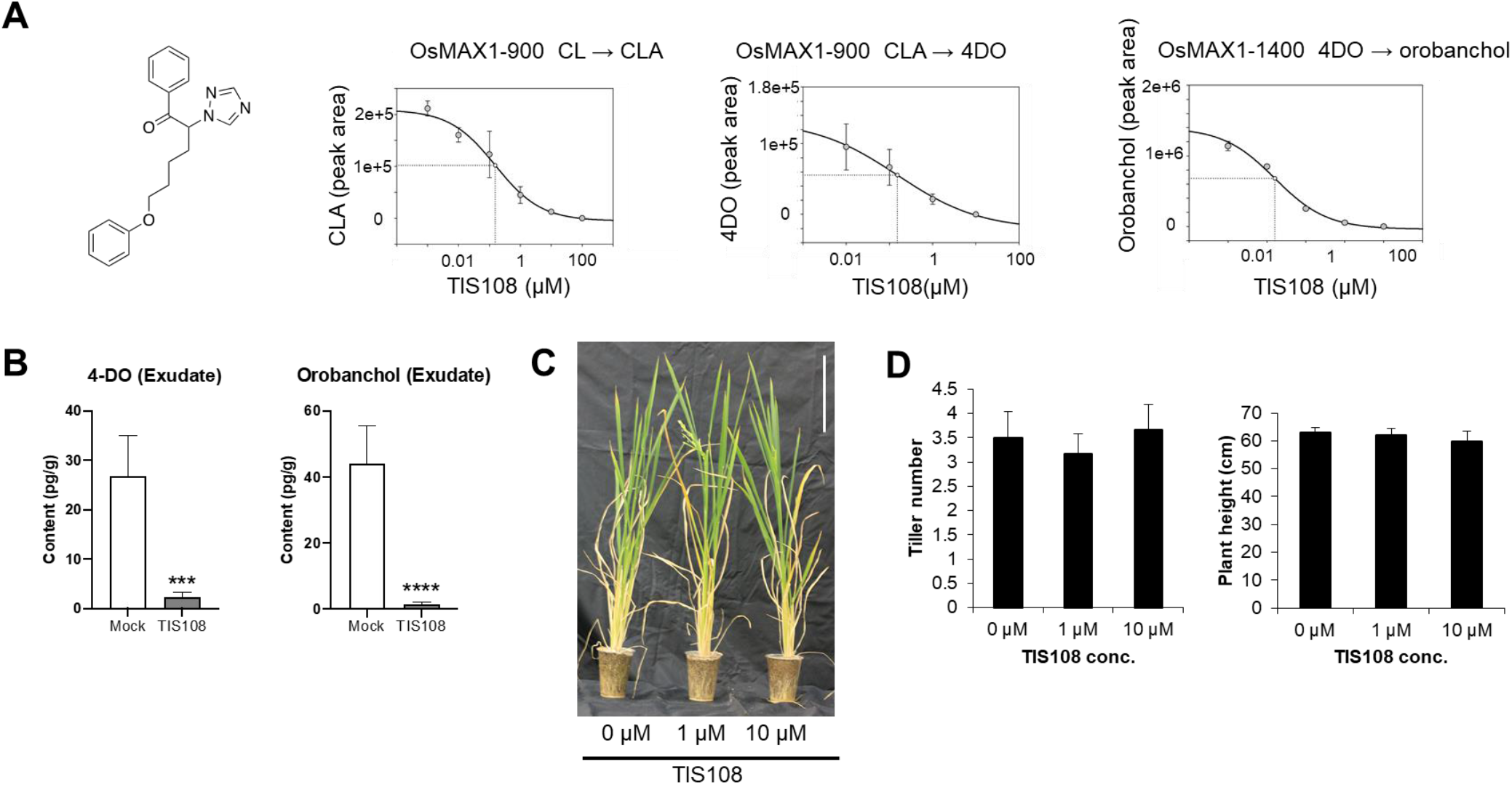
TIS108 is an OsMAX1s inhibitor. (**A**) Structure of TIS108 and inhibition of the activity of rice MAX1s by TIS108. Different substrates (carlactone, carlactonoic acid and 4-deoxyorobanchol) and concentrations of TIS108 were incubated with MAX1 containing yeast microsomes. Assay extracts and authentic standard controls were analyzed by LC-MS/MS. (**B**) TIS effect on canonical SLs, 4DO and orobanchol, in root exudates of WT grown under constant low Pi conditions. The data are presented as means ± SD of 5 biological replicates. Asterisk indicates significant difference without (Mock) and with 10 μM TIS108 treatment (TIS108) (\*\*\**P*<0.001, \*\*\*\**P*≤0.0001, Student’s t test). (**C**) Three-month-old rice plants treated with TIS108. Scale bar = 10 cm. (**D**) Tiller number and plant height of plants from (C).

To confirm the effect of TIS108 on the biosynthesis of canonical SLs *in planta* and to check its impact on plant growth and architecture, we applied the inhibitor to hydroponically grown rice seedlings under phosphate starvation. TIS108 treatment caused a significant decrease of 4DO, orobanchol, and 4-oxo-MeCLA level and an accumulation of CL+30 (Fig. 3B; Fig. S19). Seedlings of the rice *d14*-1 SL-perception mutant, which contains higher amounts of SLs due to the absence of a negative feedback regulation, showed similar responses to TIS108 treatment, i.e. a decrease of canonical SLs in roots and root exudates and an enhancement in CL+30 level (Fig. S20); confirming the impact of TIS108 on SL pattern. Importantly, the application of TIS108 to 2-week-old rice WT seedlings grown in hydroponic (Fig. S21) or soil (Fig. 3C and D) did not cause phenotypic alterations, compared to the mock. We also investigated the effect of TIS108 on rice transcriptome, using RNAseq (Data S1). None of the identified 174 upregulated and 107 downregulated differentially expressed genes (DEGs) in TIS108-treated rice (Tables S1 and S2) was related to tillering or SL biosynthesis. This result is in line with the absence of significant morphological changes upon TIS108 treatment (Table S3). Furthermore, we investigated the impact of TIS108 on the AM symbiosis. Application of this inhibitor at a 10 μM concentration to plants inoculated with the AMF *R. irregularis* led to a colonization pattern, based on *OsPT11* transcript abundance, similar to that observed with the *Os900*-KO mutants: TIS108 caused a delay in mycorrhization at 10 dpi, which was recovered at 20 dpi. However, by the end of the experiment, TIS108-treated plants showed a tendency towards reduction of *OsPT11* transcript level, compared to WT (Fig. S22). Next, we investigated whether TIS108 can be utilized for reducing *Striga* infestation. For this purpose, we exposed rice grown in *Striga*-infested soil to TIS108 at concentrations of 0, 0.0782, 0.235, and 0.782 mg/L (total amounts) over a 7-week time period. Results obtained showed a reduction of *Striga* emergence in a dose-dependent manner (Fig. 4A-E; Fig. S23-S24). We did not observe this decrease when we added the SL analog methyl-phenlactonoate 1 (MP1; Jamil et al. 2018) to the TIS108 treatment, suggesting that the lower *Striga* emergence detected with TIS108 alone is a result of lower level of germination stimulant in the root exudates. Lower infestation protected the rice plants from *Striga*-induced growth inhibition (Fig. 4A), leading to number of tillers and spikes, plant height, grain yield, and grain number similar to those of WT rice grown in *Striga*-free soil and without TIS108 treatment (Fig. 4B-E; Fig. S23). We also tested the effect of TIS108 on Indica rice and sorghum - major crops in *Striga* infested regions in Africa. Here again, we observed lower *Striga* germination inducing activity of the exudates isolated from TIS108 treated plants (Fig. S25). Overall, the application of TIS108 mimics the effect of knocking out *MAX1-900* in the *Osmax1-900* mutants (Fig. S26), with respect to the level of canonical SLs and biological activity of root exudates, suggesting that rice canonical SLs are rhizospheric signals rather than tillering-inhibitory hormones.

**FIGURE 4.**
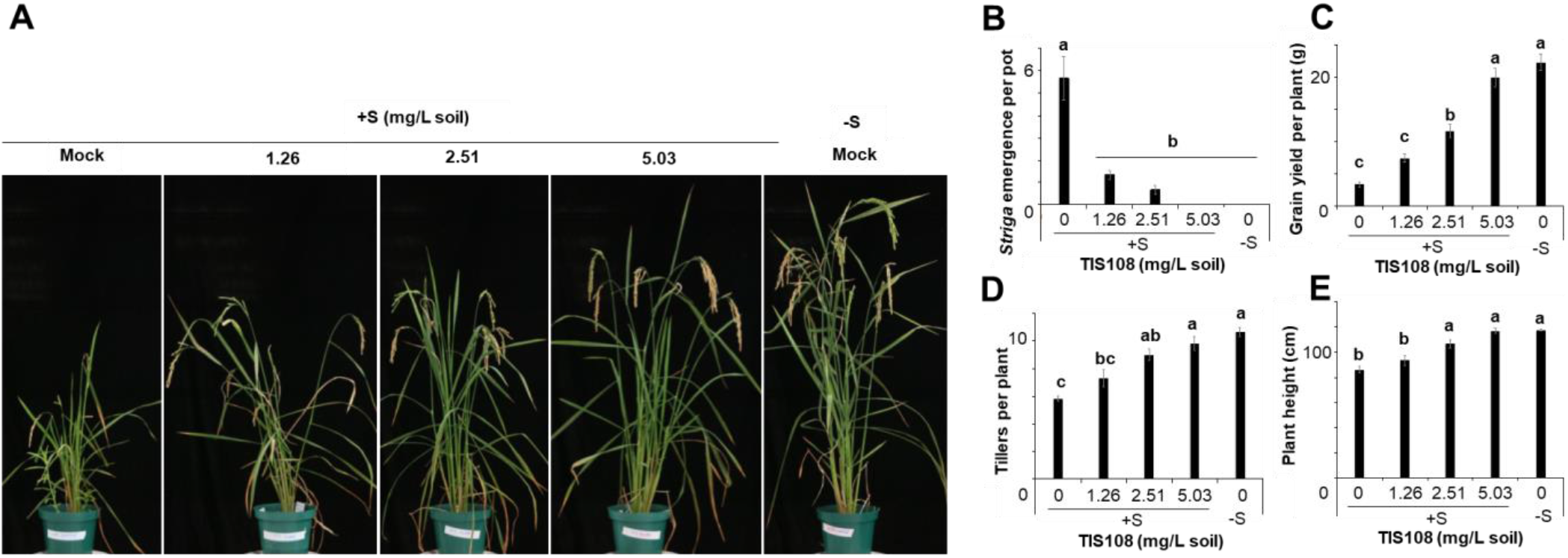
Application of TIS108 mitigates *Striga* infestation. (**A**) *Striga* emergence test in rice grown in the presence (+S) or absence (-S) of *Striga* seeds for 8 weeks. The soil was treated with 0, 10, 20 or 40 µM TIS108 once a week up to 3 weeks. Total amounts of TIS108 were 1.26 (10 µM TIS108), 2.51 (20 µM TIS108) and 5.03 (40 µM TIS108) mg/L soil. (**B**) Number of emerged *Striga* plants after 8 weeks. Grain yield (**C**), number of tillers (**D**) and plant height (**E**) were recorded at final harvesting. The data are presented as means ± SE from 6 samples. Different letters indicate statistically significant differences at *P*_0.05_.

Taken together, we employed genetic and chemical strategies to manipulate rice SL compositions, which allowed us to disentangle the biological functions of canonical and non-canonical SLs in rice. Our findings unraveled the possibility of reducing *Striga* infection by gene editing or chemical treatment without significantly affecting host’s morphology, growth and symbiotic capability. For immediate practical purpose, we estimated the effective concentration of TIS108 to be around 305 g/ha, and its application is a promising strategy alleviating the threat posed by *Striga* and other root parasitic plants to global food security.

## Author Contributions

Conceptualization, S.I., J.Y.W., J.B., T.N., T.A, and S.A.-B.;

Investigation, S.I., J.B., J.Y.W., A.Y., V.F., T.M., M.J., L.B., A.B., C.R., S.A., I.M., I.T., K.K., S.M., A.F., A.S., S.A., and N.T.;

Generation of the *Os900*-KO lines, J.B., A.F., I.H. and S.A.;

Carlactone and carlactonoic acid synthesis, A.B. and K.A.;

Phenotyping studies, J.B., J.Y.W., M.J., C.R., L.B.;

Characterization Os900-KO lines by LC-MS/MS analysis, J.Y.W. and J.B.;

Rice feeding experiments with zaxinone and 4DO, J.Y.W. and J.B.;

AMF related studies, V.F., T.M. and L.L.;

Root parasitic plant studies, J.B., J.Y.W., I.T., and M.J.;

TIS108 related LC-MS/MS analysis, S.I., J.Y.W., J.B., A.Y., and T.N.;

TIS108 Synthesis, S.I.;

Expression of SL biosynthesis genes, A.Y. and T.N.;

RNAseq analysis, S.I., S.M., and A.S.;

Resources, S.I., L.F., T.N., T.A., and S.A.-B.;

Writing – Original Draft, J.B., J. Y. W. and S.A.-B.;

Writing – Review & Editing, S.I., J.B., J. Y. W., V.F., M. J., Y.S., S.Y., L.L., M.Z., T.N., T.A., and S.A.-B.

Funding acquisition, S.I., T.N., T.A. and S.A.-B.

## Acknowledgments

We thank A. Gabar Babiker (Natinola Center for Research, Sudan) for providing *Striga* seeds and Haroon Butt for the *d17* seeds. We are grateful to Salim Sioud and Vasileios Samaras from KAUST Analytical Chemistry Corelab for their assistance in the SL identification. We also thank Tomoyasu Sato, Elle Kanbayashi and Mara Novero for technical assistance with the experiments.

