## Supplementary information for "Canonical Strigolactones Are Not the Tillering-Inhibitory Hormone but Rhizospheric Signals in Rice"

#### Material and Methods

##### Plant and fungi materials

Rice (*Oryza* sativa, cv. Nipponbare) mutant lines were generated in previous studies: *d17* (dl-2) (Butt et al. 2018) and *d10* (cv. Shiokari) (Ito et al. 2010). Their respective wild-type backgrounds were used in the experiments.

*R. irregularis* DAOM 197198 was obtained from Agronutrition, in Labège, France.

*Striga hermonthica* seeds were collected from sorghum fields during the 2012 rainy season in Sudan and were generously provided by Prof. Abdel Gabbar Babiker. *Phelipanche ramosa* seeds were kindly provided by Philippe Semier, Université de Nantes, France.

##### Generation of Os900-KO plants

Rice (*Oryza sativa* L. ssp. japonica cv. Nipponbare) *OsMAX1-900* (*Os01g0700900*/JX235697) gene was targeted using CRISPR/Cas9 guided by two gRNAs (sgRNA1: 5'-ggaagtacggccccatcttc-3', sgRNA2: 5'-aattctcctgttcatcagaa-3'), designed using the CRISPR-PLANT database *(29)*. The construction of the tRNA-gRNA-Cas9 cassette was done through Golden Gate assembly into pRGEB32 binary vector containing hygromycin resistance *(30)*. Induced Nipponbare calli, from mature seeds, were transformed with *Agrobacterium tumefaciens* EHA105 culture containing the plasmid of interest, selected and regenerated in the presence of hygromycin. After successive regeneration of shoots and roots in a Percival growth chamber (CLF Plant Climatics GmbH, model CU 36L5), the plantlets were transferred to soil and grown in a greenhouse at 28°C Day/22°C Night *(31)*.

Plant transgenicity was tested by PCR amplification of the region surrounding the two sgRNAs insertion site in the pRGEB32 vector with pRGEB32 specific primers pRGEB32 -F (5'-ccacgtgatgtgaagaagtaagataaactg-3'), and pRGEB32-R (5'-

gataggtttaagggtgatccaaattgagac-3’). The CRISPR-mediated mutations were identified by amplifying the DNA region that contains the sgRNAs target sites with the genome specific primers Os900 sg2-sg3 F (5’-gccatactggaaagtgcgg-3’) and Os900 sg2-sg3 R (3’-tagcttcaggtaaaattgcgcg-5’) and by sequencing the resulting 455bp long PCR fragment (Fig. S27).

**Hydroponic culture of rice seedlings**

Seeds from $T_3$ homozygous plants were first surface sterilized in 2% sodium hypochlorite (v/v) with 0.01% Tween-20 for 20 min under gentle agitation before being rinsed generously with sterile water and germinated overnight in the dark (30 °C). The pre-germinated seeds were then transferred to round Petri dishes containing two sheets of sterile Whatman filter paper and 5 mL of half-strength Murashige and Skoog (MS) (PhytoTechnology Laboratories, Cat. No. M519) solution (pH 5.7) and incubated for another 2 days in dark at 30 °C. Finally, 15 mL of modified Hoagland nutrient solution adjusted at pH 5.8 (0.4 mM $K_2HPO_3$*$3H_2O$, 5.6 mM $NH_4NO_3$, 0.8 mM $MgSO_4$*$7H_2O$, 0.8 mM $K_2SO_4$, 0.18 mM $FeSO_4$*$7H_2O$, 0.18 mM $Na_2EDTA$*$H_2O$, 1.6 mM $CaCl_2$, 0.8 mM $KNO_3$, micronutrients (0.023 mM $H_3BO_3$, 4.5 µM $MnCl_2$ *$4H_2O$, 0.3 µM $CuSO_4$*$5H_2O$, 1.5 µM $ZnCl_2$, 0.1 µM $Na_2MoO_4$*$2H_2O$)) was added, and seedlings were incubated in a Percival for 5 days (day/night temperature of 28/22 °C and a 12 h photoperiod, 200 µmol photons $m^{-2}$ $s^{-1}$.

The set-up of the hydroponic culture consisted of 50 mL black tubes with a perforated cap containing in its center a 1.5 mL bottomless Eppendorf tube, to which the 1-week-old seedlings were transferred into. The nutrient solution provided with or without 0.4 mM $K_2HPO_3$*$3H_2O$ resulted in the +Pi and -Pi conditions, respectively. -Pi conditions were achieved by feeding the seedling with +Pi for two weeks, followed by one week of -Pi treatment prior further analysis (seedling phenotyping). The solutions were changed every three days.

For low Pi condition, the same procedure was performed but with replacing the half-strength MS solution by 5 mL of modified Hoagland nutrient solution adjusted at pH 5.8, containing 0.004 mM $K_2HPO_3*3H_2O$ (low Pi), right after the sterilization step. Plants were kept for three weeks in low Pi solution.

#### Phenotyping in pots and rhizotron

For phenotyping of *Os900* mutants, the seedlings were transferred into pots filled with soil containing half-strength modified Hoagland nutrient solution. The nutrient solution consisted of 5.6 mM $NH_4NO_3$, 0.8 mM $MgSO_4{\cdot}7H_2O$, 0.8 mM $K_2SO_4$, 0.18 mM $FeSO_4{\cdot}7H_2O$, 0.18 mM $Na_2EDTA{\cdot}2H_2O$, 1.6 mM $CaCl_2{\cdot}2H_2O$, 0.8 mM $KNO_3$, 0.023 mM $H_3BO_3$, 0.0045 mM $MnCl_2{\cdot}4H_2O$, 0.0003 mM $CuSO_4{\cdot}5H_2O$, 0.0015 mM $ZnCl_2$, 0.0001 mM $Na_2MoO_4{\cdot}2H_2O$, and with or without 0.4 mM $K_2HPO_4{\cdot}2H_2O$, resulting in the +Pi and -Pi conditions, respectively. The pH of the solution was adjusted to 5.8, and the solution was applied every third day. On day 56, phenotypic data were recorded. The plants were grown in a greenhouse from February to April 2020, in Thuwal (Saudi Arabia).

For observing the *Os900* mutant root phenotypes in the rhizotron system (48 cm × 24 cm × 5 cm), 3 days-old seedlings were grown in soil with Hoagland nutrient solution containing 0.4 mM $K_2HPO_4{\cdot}2H_2O$ (+Pi) for 2 weeks. The solution was changed every other day with fresh nutrient solution. Root length, angle and surface area were analyzed with the ImageJ software.

#### Qualitative and quantitative analysis of SLs in root exudates, root tissues, and shoot base

Analysis of SLs in rice root exudates and root tissues was performed following the protocol described by Wang et al. (Wang et al. 2019). Briefly, collected 50 mL root exudates of two seedlings grown together in one tube, and spiked with 0.672 ng of $D_6$–5DS, were brought on a $C_{18}$-Fast Reversed-Phase SPE column (500 mg/3 mL; GracePure™) preconditioned with 3 mL of methanol and 3 mL of water. After washing with 3 mL of water, SLs were eluted with 5 mL of acetone. The SLs fraction was concentrated to SL aqueous solution (~1 mL), followed by 1 mL of ethyl acetate extraction. 750 µL of SL enriched organic phase was dried under vacuum. For root tissues analysis, around 25 mg of lyophilized and grinded root tissues, spiked with 0.672 ng of

$D_6$–5DS, were extracted twice with 2 mL of ethyl acetate in an ultrasound bath (Branson 3510 ultrasonic bath) for 15 min, followed by centrifugation for 8 min at 3800 rpm at 4 °C. The two supernatants were combined and dried under vacuum. The residue was dissolved in 50 µL of ethyl acetate and 2 mL of hexane, followed by a Silica Cartridges SPE column (500 mg/3 mL; HyperSep™) purification. After washing with 3 ml of hexane, SLs was eluted in 3 ml of ethyl acetate and evaporated to dryness under vacuum. The same procedure was used for shoot base (shoot-root junction) tissue extraction, except for the tissue powder preparation: 12 fresh shoot bases were pulled together and manually grinded, while preserved liquid nitrogen. The entire sample was used during the extraction. The final extract was re-dissolved in 100 µL of acetonitrile: water (25:75, v:v) and filtered through a 0.22 µm filter for LC-MS/MS analysis.

SLs were identified by using UHPLC-Orbitrap ID-X Tribrid Mass Spectrometer (Thermo Scientific™ Altis™) with a heated-electrospray ionization source. Chromatographic separation was achieved on the Hypersil GOLD $C_{18}$ Selectivity HPLC Columns (150 × 4.6 mm; 3 µm; Thermo Scientific™) with mobile phases consisting of water (A) and acetonitrile (B), both containing 0.1% formic acid, and the following linear gradient (flow rate, 0.5 mL/min): 0–15 min, 25%–100 % B, followed by washing with 100 % B and equilibration with 25 % B for 3 min. The injection volume was 10 µL, and the column temperature was maintained at 30 °C for each run. The MS conditions were as follows: positive mode, ion source of H-ESI, spray voltage of 3500V, sheath gas flow rate of 60 arbitrary units, auxiliary gas flow rate of 15 arbitrary units, sweep gas flow rate of 2 arbitrary units, ion transfer tube temperature of 350 °C, vaporizer temperature of 400 °C , S-lens RF level of 60, resolution of 120000 for MS; stepped HCD collision energies of 10, 20, 30, 40, 50% and resolution of 30000 for MS/MS. The mass accuracy (accurate mass ± 5 ppm mass tolerance) of identified compounds and their MS spectra (accurate mass ± 5 ppm mass tolerance) were acquired using Xcalibur software version 4.1.

SLs were quantified by LC-MS/MS using a HPLC-triple quadrupole/linear ion trap instrument (QTRAP5500; AB Sciex) and UHPLC- Triple-Stage Quadrupole Mass Spectrometer (Thermo Scientific™ Altis™). Chromatographic separation was achieved on a ZORBAX Eclipse plus $C_{18}$ column (150 × 2.1 mm; 3.5 µm; Agilent) with mobile phases consisting of water:acetonitrile (95:5, v:v; A) and acetonitrile (B), both containing 0.1% formic acid, and the following linear gradient (flow rate, 0.5 mL/min): 0–15 min, 25%–100 % B, followed by washing with 100 % B and equilibration with 25 % B for 3 min. The injection volume was 10 μL, and the column temperature was maintained at 35 °C for each run. The MS parameters of QTRAP5500 were as follows: positive ion mode, ion source of turbo spray, ion spray voltage of 5500 V, curtain gas of 40 psi, collision gas of medium, gas 1 of 60 psi, gas 2 of 50 psi, turbo gas temperature of 400 °C, declustering potential of 60 V, entrance potential of 10 V, collision energy of 16 eV, collision cell exit potential of 10 V. The MS parameters of QTRAP5500 were as follows: positive ion mode, ion source of turbo spray, ion spray voltage of 5500 V, curtain gas of 40 psi, collision gas of medium, gas 1 of 60 psi, gas 2 of 50 psi, turbo gas temperature of 400 °C, de-clustering potential of 60 V, entrance potential of 10 V, collision energy of 16 eV, collision cell exit potential of 10 V. The MS parameters of Thermo Scientific™ Altis™ were as follows: positive ion mode, ion source of H-ESI, ion spray voltage of 5000 V, sheath gas of 40 arbitrary units, aux gas of 15 arbitrary units, sweep gas of 20 arbitrary units, ion transfer tube gas temperature of 350 °C, vaporizer temperature of 350 °C, collision energy of 17 eV, CID gas of 2 mTorr, and full width at half maximum (FWHM) 0.2 Da of Q1/Q3 mass. The characteristic Multiple Reaction Monitoring (MRM) transitions (precursor ion → product ion) were 331.15→216.0, 331.15→234.1, 331.15→97.02 for 4-deoxyorobanchol; 347.14→329.14, 347.14→233.12, 347.14→ 205.12, 347.14→97.02 for orobanchol; 361.16→ 247.12, 361.16→177.05, 361.16→208.07, 361.16→97.02 for 4-oxo-MeCLA isomer; 333.17→219.2, 333.17→173.2, 333.17→201.2, 333.17→97.02 for putative 4-oxo-hydroxyl-CL (CL+30); 337.19→222.15, 337.19→240.16, 337.19→97.02 for $D_6$-5-deoxystrigol.

**Feeding with $^{13}$C-labled CL**

$^{13}$C-CL was prepared by following the protocol described by Bruno et al. 2016 (Bruno and Al-Babili 2016). Briefly, the *OsCCD8* cDNA – controlled by an arabinose-inducible promoter - was expressed as thioredoxin-fusion in BL21 Rosetta *E.coli* cells. A single colony of the transformed *E.coli* was cultured overnight, from which 0.5 mL was inoculated into 50 mL media and grown at 28 °C. When the $OD_{600nm}$=0.5, we induced the protein production with 0.2% (w/v) arabinose and incubated under agitation for four hours at 28 °C. The harvested cells (centrifugation) were re-suspended in lysis buffer (sodium phosphate buffer pH8 containing 1% Triton X-100 and 10 mM of dithiothreitol, Lysozyme (1mg/ml)) and incubated on ice for 30 min. The crude lysate was then sonicated and centrifuged at 12000 rpm and 4 °C for 10 min. The protein was collected (supernatant) to be used for the *in vitro* incubation with $^{13}C$ labelled 9-cis-β-apo-10-carotenal, from Buchem B. V. (Apeldoorn, Netherlands). The substrate ($^{13}C$ labelled 9-cis-β-apo-10-carotenal) was quantified spectrophotometrically, dried and re-suspended in ethanolic detergent mixture 0.4% (v/v) Triton X-100. The mixture was then dried using a vacuum centrifuge to produce a carotenoid-containing gel, which was re-suspended in incubation buffer (2 mM tris 2-carboxyethylphosphine, 0.4 mM $FeSO_4$ and 2 mg/ml catalase (Sigma, Deisenhofen, Germany) in 200 mM Hepes/NaOH, pH 8). OsCCD8 crude cell lysate, i.e. 50 µl of the soluble fraction of overexpressing cells, was added to the assay, and the whole mix incubated 4hrs under shaking at 140 rpm at 28°C in dark. The reaction was stopped by adding two volumes of acetone, and the lipophilic compounds were separated by partition extraction with petroleum ether: diethyl ether 1: 4 (v / v), dried, and resuspended in methanol for HPLC analysis.

$^{13}C$-carlactone was preparatively purified using a YMC-Pack C30-reversed phase column (250 × 4.6 mm i.d., 5 µm) in Agilent 1260 HPLC. The following separation systems were used. The column was developed at a flow-rate of 1 ml/min with a gradient from 100% B to 80% B (MeOH:water:tert-butylmethyl ether (30: 10: 1, v/v/ v)) within 15 min, then to 100% A (methanol:tert-butylmethyl ether (1: 1, v/v)) within 0.5 min and finally to 100% A and a flow-rate of 2 ml/min within 0.5 min, maintaining the final conditions for another 14 min. The collected fractions were dried under nitrogen gas, dissolved in dichloromethane and keep at -80 °C.

Around 20 ng $^{13}C$-CL was fed to two-week old *Os900* rice seedlings for 6 h, and then 500 mL root exudates were collected for LC-MS/MS analysis.

**Gene expression analysis**

For transcript analysis, total RNA was extracted from rice roots using the Direct-zol RNA Miniprep Plus kit (Zymo research, #R2071), according to the manufacturer's instructions. cDNA was synthetized from 2 µg of total RNA using iScript cDNA Synthesis Kit (BIO-RAD Laboratories, Inc, 2000 Alfred Nobel Drive, Hercules, CA; USA) according to the instructions in the user manual. qRT-PCR was performed using SYBR Green Master Mix (Applied Biosystems), which was diluted before use. Each PCR reaction was carried out in a total volume of 10 µL containing 4 µL diluted cDNA (the resulting cDNA was diluted 1:10 in $H_2O$), 5 µL 2X diluted SYBR Green Reaction Mix, and 0.5 µL of each primer (2 µM working stock). All reactions were performed on a 384-well plate in a CFX384 Touch Real-Time PCR Detection System (Bio-Rad) as follows: 95 °C for 90 sec, 40 cycles of 95 °C for 15 sec, 60 °C for 30 sec. All reactions were performed on at least three biological and three technical replicates, including a water control to exclude potential unspecific amplification. The $2^{-\Delta\Delta C_T}$ method was used to calculate the relative gene expression levels (Livak and Schmittgen 2001) and rice Ubiquitin (*OsUBQ*) gene was used as the internal control to normalize target gene expression (see Table S4 for primer sequences).

For *OsPT11* gene expression analysis (AMF marker), total RNA was extracted from rice roots using the Plant RNeasy Kit (Qiagen), according to the manufacturer's instructions. Samples were treated with TURBO™ DNase (Ambion) according to the manufacturer's instructions. The RNA samples were routinely checked for DNA contamination by means of PCR analysis, using primers for *OsRubQ1* (Guimil et al. 2005). For single-strand cDNA synthesis about 1000 ng of total RNA was denatured at 65 °C for 5 min and then reverse-transcribed at 25 °C for 10 min, 42 °C for 50 min, and 70 °C for 15 min. The reaction was carried out in a final volume of 20 µL containing 10 µM random primers, 0.5 mM dNTPs, 4 µL 5X buffer, 2 µL 0.1 M DTT and 1 µL Super-Script II (Invitrogen). Quantitative RT-PCR (qRT-PCR) was performed using a Rotor-Gene Q 5plex HRM Platform (Qiagen). Each PCR reaction was carried out in a total volume of 15 µL containing 2 µL diluted cDNA (about 10 ng), 7.5 µL 2X SYBR Green Reaction Mix, and 2.75 µL of each primer (3 µM). The following PCR program was used: 95°C for 90 sec, 40 cycles of 95 °C for 15 sec, 60 °C for 30 sec. A melting curve (80 steps with a heating rate of 0.5 °C per 10 sec and a continuous fluorescence measurement) was recorded at the end of each run to exclude the generation of non-specific PCR products. All reactions were performed on at least three biological and three technical replicates. Baseline range and take off values were automatically calculated using Rotor-Gene Q 5plex software. Transcript level of *OsPT11* (Guimil et al. 2005) were normalized using *OsRubQ1* housekeeping gene (Guimil et al. 2005). Only take off values leading to a Ct mean with a standard deviation below of 0.5 were considered.

**Exogenous applications of 4DO and zaxinone**

For investigating the effect of 4DO (purchased from OlChemIm; Czech Republic) on *d1*7 mutant, 1-week-old seedlings were grown hydroponically in half-strength Hoagland nutrient solution containing 0.4 mM $K_2HPO_4.2H_2O$ (+Pi) with different concentration of 4-DO for 2 weeks. 1000 nM 4DO (dissolved in 0.1% acetone) was prepared as stock solution, and using serial dilution to make 100 nM, 10 nM, and 1 nM 4DO solution respectively. 1000 nM *rac*-GR24 was used as positive control and the same amount of acetone was adjusted in each group. The solution was changed twice per week, adding the chemical at each renewal.

For investigating the effect of zaxinone (customized synthesis from Buchem B.V.; Apeldoorn, The Netherlands.) on different genotypes, 1 week-old seedlings were grown hydroponically in half-strength Hoagland nutrient solution containing 0.4 mM K2HPO4.2H2O (+Pi), 2.5 µM zaxinone (dissolved in 0.1% acetone), 1 µM rac-GR24 (purchased from StrigoLab; Turin, Italy), or the corresponding volume of the solvent (mock; acetone) for 2 weeks. The solution was changed twice per week, adding the chemical at each renewal.

**Plant material and growth conditions for *R. irregularis* root colonization**

Seeds of WT plants and *Osmax1* independent lines (*Os900-32* and *-34*) were germinated in pots containing sand and incubated for 10 days in a growth chamber under a 14 h light (23°C)/10 h dark (21°C). Plants used for mycorrhization were inoculated with ~ 1000 sterile spores of *R. irregularis* DAOM 197198 (Agronutrition, Labège, France). A set of WT mycorrhizal plants were treated with TIS108 (10 µM), once per week, by applying the compound once a week directly in the nutrient solution. Non-mycorrhizal and mycorrhizal plants were grown in sterile quartz sand and watered with a modified Long-Ashton (LA) solution containing 3.2 μM $Na_2HPO_4 \cdot 12\ H_2O$ (Hewitt, 1966). Mycorrhizal level was monitored during a time-course experiment from 10, 20 to 35 days-post-inoculation (dpi). In the last time point (35 dpi) wild-type and *Os900* mycorrhizal roots were stained with 0.1 % cotton blue in lactic acid and the estimation of mycorrhizal parameters was performed by Trouvelot method (Trouvelot, Kough, and Gianinazzi-Pearson 1986). Four parameters were considered: F%, percentage of segments showing internal colonization (frequency of mycorrhization); M%, average percentage of colonization of root segments (intensity of mycorrhization); a%, percentage of arbuscules within infected areas; A%. For the phenotype evaluation of non-mycorrhizal and mycorrhizal plants, we considered the following parameters: crown root (CR) number, root and shoot length, leaves number and root and shoot fresh weight.

**Gene expression analysis of mycorrhizal plants**

Total RNA was extracted from rice roots using the Plant RNeasy Kit (Qiagen), according to the manufacturer's instructions. Samples were treated with TURBO™ DNase (Ambion) according to the manufacturer's instructions. The RNA samples were routinely checked for DNA contamination by means of PCR analysis, using primers for *OsRubQ1* (Guimil et al. 2005). For single-strand cDNA synthesis about 1000 ng of total RNA was denatured at 65°C for 5 min and then reverse transcribed at 25°C for 10 min, 42°C for 50 min, and 70°C for 15 min. The reaction was carried out in a final volume of 20 μL containing 10 μM random primers, 0.5 mM dNTPs, 4 μL 5X buffer, 2 μL 0.1 M DTT and 1 μL Super-Script II (Invitrogen). Quantitative RT-PCR (qRT-PCR) was performed using a Rotor-Gene Q 5plex HRM Platform (Qiagen). Each PCR reaction was carried out in a total volume of 15 μL containing 2 μL diluted cDNA (about 10 ng), 7.5 μL 2X SYBR Green Reaction Mix, and 2.75 μL of each primer (3 μM). The following PCR program was used: 95°C for 90 sec, 40 cycles of 95°C for 15 sec, 60°C for 30 sec. A melting curve (80 steps with a heating rate of 0.5°C per 10 sec and a continuous fluorescence measurement) was recorded at the end of each run to exclude the generation of non-specific PCR products. All reactions were performed on at least three biological and three technical replicates.

Baseline range and take off values were automatically calculated using Rotor-Gene Q 5plex software. Transcript level of OsPT11 (Guimil et al. 2005) were normalized using *OsRubQ1* housekeeping gene (Guimil et al. 2005). Only take off values leading to a Ct mean with a standard deviation below of 0.5 were considered. Statistical tests were carried out through one-way analysis of variance (one-way ANOVA) and Tukey's post hoc test, using a probability level of $p<0.05$. All statistical elaborations were performed using PAST statistical (version 2.16; (Hammer, Harper, and Ryan 2001).

**Spore germination assay**

Spores were sterilized in a solution of streptomycin sulphate (0.03% W/V) and chloramine T (3% W/V) and germinated in 100 µl of *max1-32* and *max1-34* root exudates at $10^{-9}$ M (CL+30 content was determined by $D_6$-5DS), GR24 $10^{-9}$ M, or a solution of water/acetone (0.08%). For each treatment, 96 sterilized spores were placed individually in the wells of a multi-well plate and treated with freshly prepared solutions at the beginning of the experiment. Spores were germinated in the dark at 30°C and the germination rate was evaluated after 3 days.

***Striga hermonthica* and *Phelipanche ramosa* germination bioassays**

*Striga hermonthica* and *Phelipanche ramosa* seeds were first sterilized and pre-conditioned as described before (Matusova et al. 2005). The pre-conditioned seeds were spread on a 9 mm-diameter Whatman filter paper disc. Each disc contains from 40 up to 100 seeds. Then, 5 discs were moved to a plastic petri dish, where each disc was used as a technical replicate. Eluted in acetone (the same procedure was followed for root exudates' extracts to be used for bioassay experiments, omitting the addition of labelled SL as the internal standard), the root exudates were added to water (300 µL of sample in 300 µL of $H_2O$) to allow acetone evaporation under vacuum-centrifugation while minimizing samples degradation, and was applied on each disc containing pre-germinated seeds. Corresponding volumes of *rac*-GR24 (purchased from StrigoLab; Turin, Italy) and sterile MilliQ water were included as positive and negative control, respectively; knowing that seeds would germinate only in presence of SL-like compounds.

The petri dishes were sealed with parafilm, enfolded with aluminum foil, and incubated for 24 h at 30°C for *Striga* and 3 days at 28°C for *Phelipanche*. The germinated and non-germinated *Striga* seeds containing disks were photographed using a Leica LED3000 R binocular microscope, adjusted to 50% medium light, mounted with a CCD camera (Leica Microsystems). The germination percentage of the acquired images was assessed by the seed counter software SeedQuant (Braguy et al. 2021).

**Heterologous expression of SL biosynthetic genes in yeast**

Heterologous expressions of *Os900* and *Os1400* in yeast (*Saccharomyces cerevisiae*) was carried out as described previously (Yoneyama et al. 2018). Yeast microsomes (approximately 100 µg proteins/100 µl) were incubated with 0.001–1000 µM TIS108 (1 µM substrate *rac*-CL and 500 µM NADPH) at 28 °C for 1 min. The reaction was stopped with the addition of 1 mL ethyl acetate. The ethyl acetate phase was dried with anhydrous sodium sulfate and then subjected to LC-MS/MS analysis as described previously (Abe et al. 2014; Yoneyama et al. 2018). The products CLA, 4DO, and orobanchol were quantified from peak areas in the transitions of *m/z* 331.1 to 113.0 in negative mode, *m/z* 331.1 to 97.0 in positive mode, and *m/z* 347.0 to 97.0 in positive mode, respectively. The $IC_{50}$ values were obtained using triplicate samples and calculated with SigmaPlot software (Systat Software, Inc., CA, USA).

**Phenotyping TIS108-treated rice**

Rice (*Oryza sativa* L. 'IAC-165') cultivation and chemical treatment was performed for 2 weeks as previously described (Ito et al. 2010). For prolonged cultivation, 2-week-old rice seedlings were planted in vermiculite and grown in the same conditions, TIS108 (100 mM) was diluted 1:10,000 from an acetone-dissolved stock solution in a hydroponic culture solution *(3)* and 100 mL was applied to the pots once a week for 10 weeks. Acetone was used as the control treatment.

**RNA sequencing**

Rice was grown in the same conditions as describe for the SL analysis. Fourteen-day-old seedlings were transferred to a brown vial with or without 10 µM TIS108 for 1 day. Roots were harvested and total RNA was extracted using an RNA purification reagent (Invitrogen, USA). The quality of total RNA was evaluated using an Agilent 2100 Bioanalyzer (Agilent, USA). Each library was prepared with NEB Next Ultra II RNA Prep Kit for Illumina according to the manufacturer's protocol (NEB, USA). The quality of each library was assessed using an Agilent 2100 Bioanalyzer and then sequenced using an Illumina NextSeq Sequencer (single-end sequencing, 75 bp). The data sets from this sequence have been deposited in the DDBJ database (accession number: DRA009250). Total reads were mapped to the rice transcripts using bowtie (Langmead et al. 2009). Differential gene expression was examined using DESeq2 (Love, Huber, and Anders 2014) and established by fold change ($\log_2$ Ratio) and false discovery rate (FDR) ($\log_2$ Ratio ≧ 1 and FDR ≦ 0.05).

##### ***Striga hermonthica*** **pot test**

Seeds of *S. hermonthica* (3 mg) were sown in plastic pots (70 mm i.d. and 84 mm in height) containing 150 mL soil (Bonsol No.2 (Sumitomo Chemical, Osaka, Japan) and river sand = 1:1, v/v) and 60 mL distilled water. Seeds in the pots were conditioned in the dark at 30 °C for 7 days. On the 8$^{th}$ day, one 5-day-old rice seedling (*Oryza sativa* L. 'IAC-165') was then planted in each *S. hermonthica* conditioned pot and grown in growth chamber (NK System, Tokyo, Japan) at 30 °C under LED light (500 µmol $m^{-2}$ $s^{-1}$) with long-day conditions (16 h light/8 h dark). Each pot was treated with 50 mL 0.1, 0.3 or 1 µM TIS108 and MP1 (Kountche et al. 2019; Jamil et al. 2018; Kountche et al. 2018) once a week for 7 weeks. The total amount of TIS108 applied was 0.0782, 0.235, and 0.782 mg/L soil. After 8 weeks, the number of *S. hermonthica* plants that emerged from each pot was counted and photographically recorded.

For *S. hermonthica* pot test, seeds of *S. hermonthica* (20 mg) were thoroughly mixed in 1.5 L sand and soil mixture (1:1) and added to 3-L perforated plastic pot containing 0.5 L clean soil in the bottom. The pots were kept in greenhouse-controlled conditions at 30°C in moist conditions for 10 days to precondition the *Striga* seeds. On the 11th day, five-day-old rice seedlings (Oryza sativa L. 'IAC-165') were planted in the middle of each pot. After 3 days, each pot was treated with 250 mL TIS108 at 10, 20 or 40 µM as an irrigation application. The compound was applied once a week for 3 weeks. The total amount of TIS108 applied was 0.84, 1.68, and 3.35 mg/L soil. The untreated pots with (+S) and without (-S) *Striga* were included as control treatments. The number of emerged *Striga* plants was recorded from each pot up to 8 weeks after initial *Striga* emergence. Rice growth, yield, and yield components were observed at the time of final harvesting.

**Statistical analysis**

Data are represented as mean and their variations as standard deviation. The statistical significance was determined by one-way analysis of variance (one-way ANOVA) and Tukey's multiple comparison test, using a probability level of $p<0.05$. All statistical elaborations were performed using GraphPad Prism 8 for Mac OS, version 8.3.0.

**Data availability**. All data generated or analyzed during this study are included in this published article and its supplementary information files. The RNA-seq data are deposited in the DDBJ Read Archive under accession no. DEA009250. RPKM values of all genes in the RNA-seq analysis are presented in data S1. All data needed to evaluate the conclusions in the paper are available upon request.

### Supplementary figures

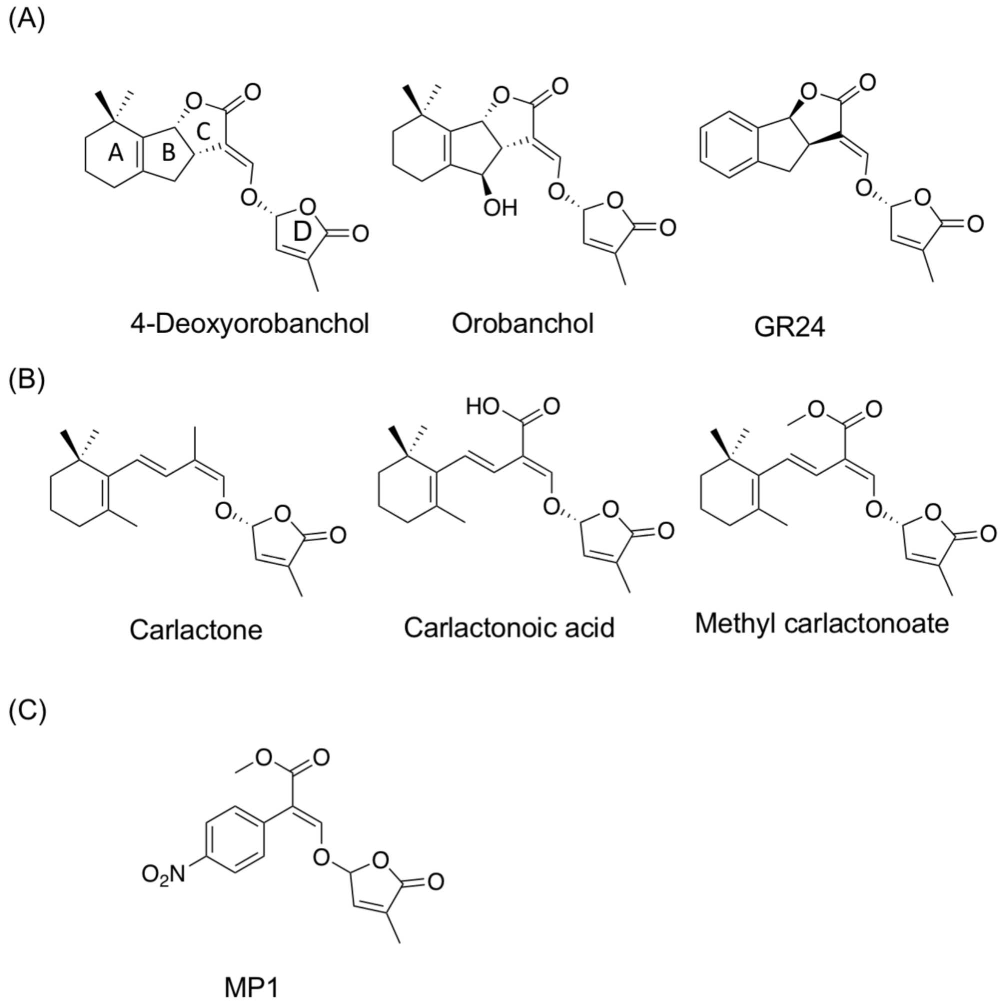


Fig. S1 Structures of canonical SLs (**A**), non-canonical SLs (**B**) and SL mimics used in this study (**C**). Abbreviations: MP1, Methyl-Phenlactonoate 1

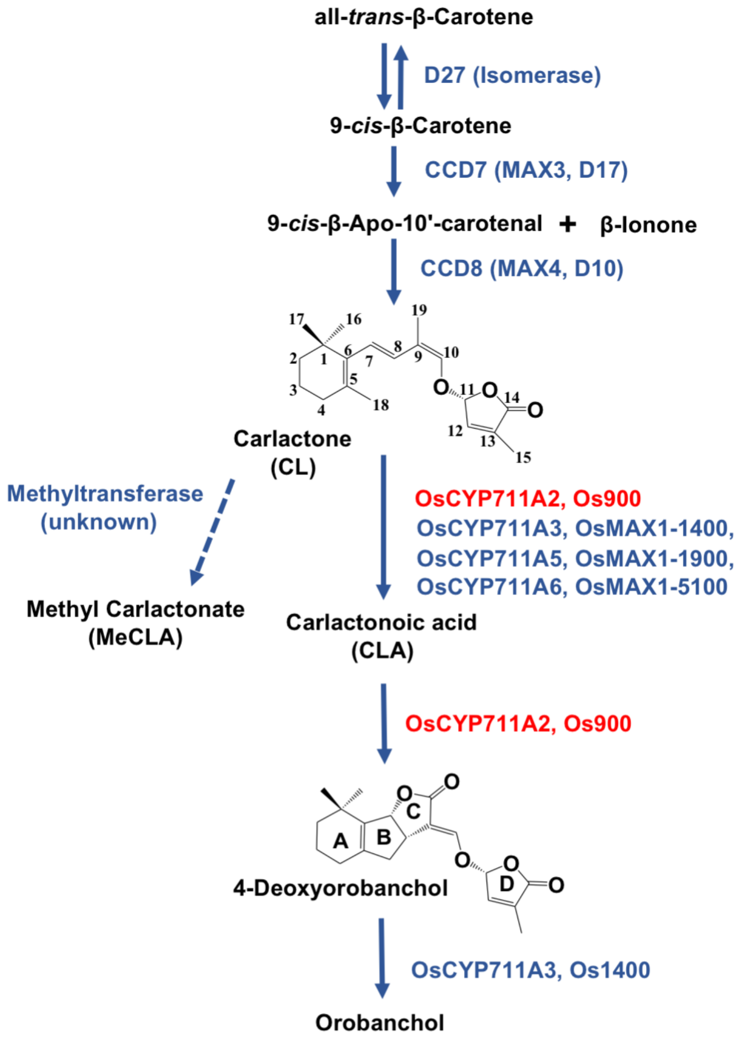


Fig. S2 SL biosynthetic pathway in rice. Abbreviations: D27, Dwarf27; CCD, Carotenoid Cleavage Dioxygenase; MAX1, More Axillary Growth 1; CYP, Cytochrome P450

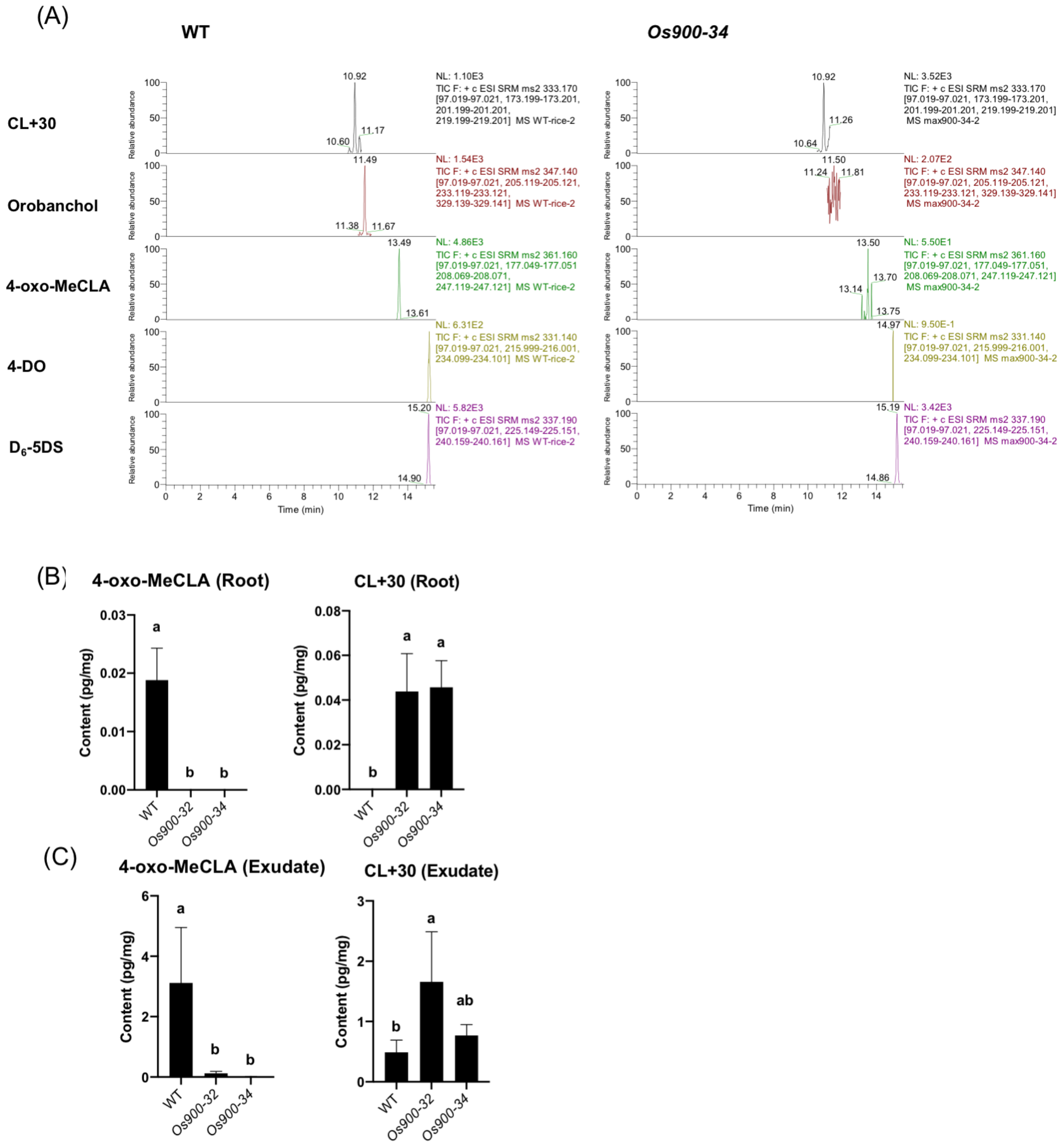


Fig. S3 Chromatograms of the different SLs. Multiple reaction monitoring (MRM) was used to identify endogenous levels of SLs in WT and *Os900-34* in root exudates (**A**) of plants grown under low Pi grown conditions. Analysis of putative non-canonical SLs, tentative 4-oxo-methyl-carlactonoate (4-oxo-MeCLA; structure proposed by Yoneyama et al., 2018) and CL+30, in root tissues (**B**) and root exudates (**C**) of *Os900*-KO lines, compared to WT, both grown under constant low Pi conditions. The data are presented as means ± SD from five biological samples. Means not sharing a letter in common differ significantly at $P_{0.05}$. Abbreviations: CL, carlactone; 4-oxo-MeCLA, 4-oxo-methyl-carlactonoate; 4-DO, 4-deoxyorobanchol; 5-DS, 5-deoxystrigol; WT, wild-type.

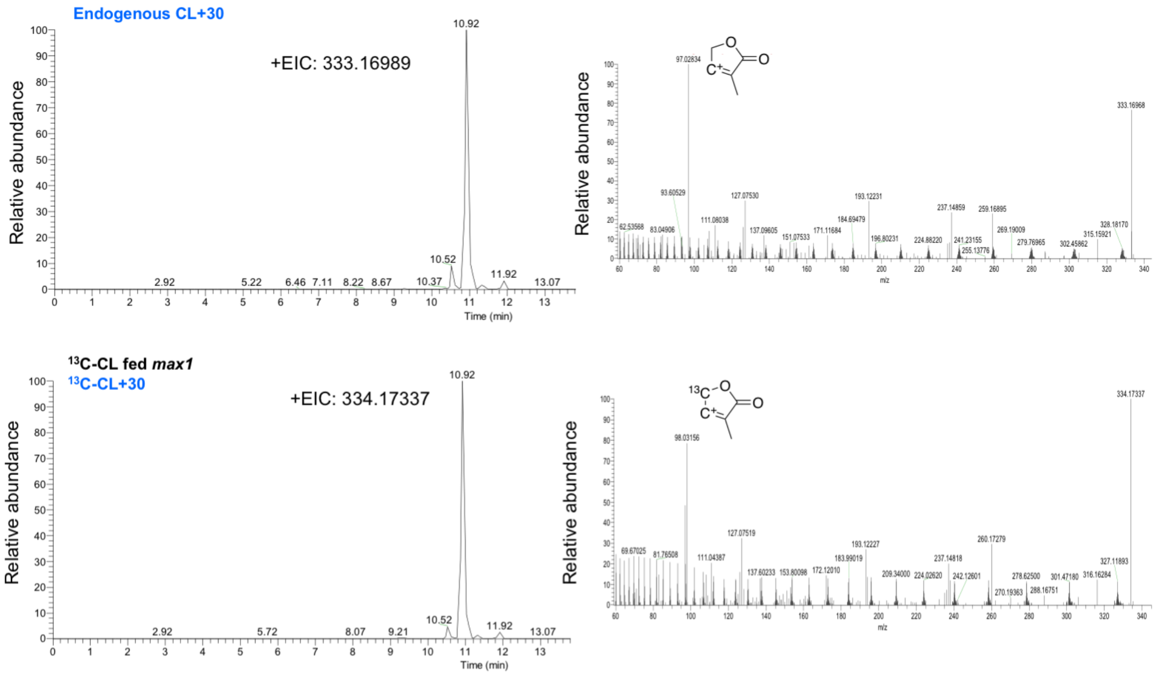


Fig. S4 Conversion of $^{13}$C-labbeled CL to CL+30. Identification of endogenous CL+30 (tentatively 4-oxo-hydroxyl-CL) (Retention time: 10.89). Product ion spectra derived from the precursor ion (m/z 333.16968 $[M+H]^+$ in positive mode) with characterized D-ring at m/z 333.16968 > 97.02834. [$^{13}$C]-CL+30 was characterized with ions pairs at m/z 334.17337>98.03156. The proposed structures of fragments are inserted.

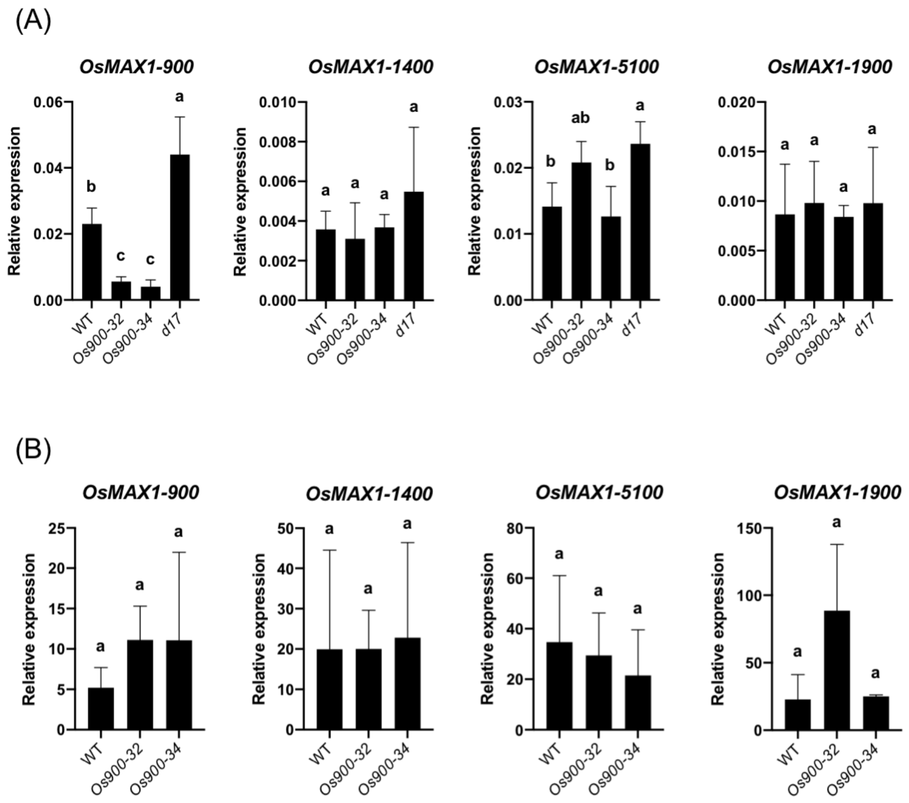


Fig. S5 Transcript analysis of the SL biosynthetic genes under (**A**) normal condition and (**B**) phosphate deficiency. The data are presented as means ± SD from three biological samples. Means not sharing a letter in common differ significantly at $P_{0.05}$. Abbreviations: *MAX1, More Axillary Growth 1*

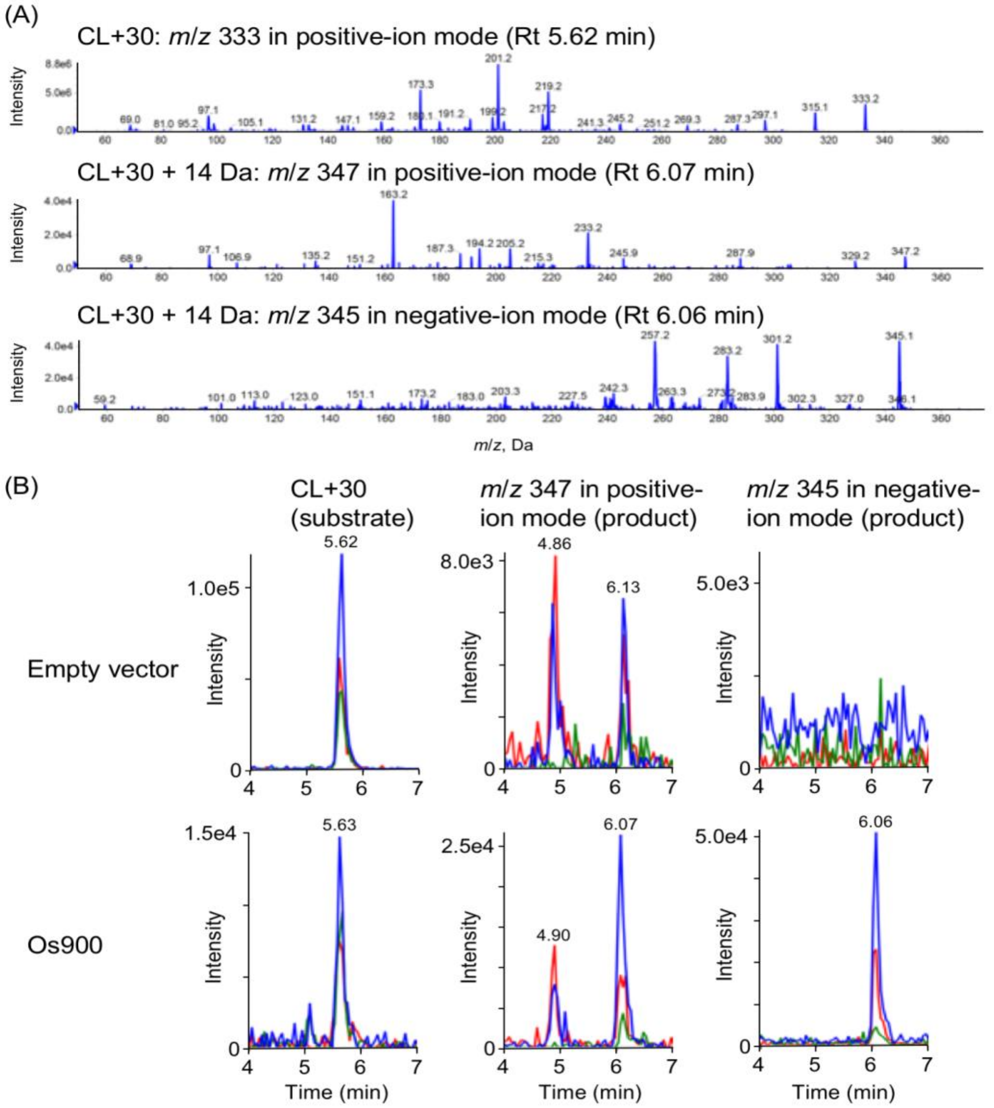


Fig. S6 Enzymatic conversion of CL+30 containing fraction to CL+30+14 Da by Os900 (**A**) Product ion spectra derived from the precursor ion (*m*/*z* 333 $[M+H]^+$ in positive mode) of substrate CL+30 and the precursor ion (*m*/*z* 347 $[M+H]^+$ in positive mode and *m*/*z* 345 $[M-H]^-$ in negative mode) of CL+30 + 14 Da produced by Os900 are shown. (**B**) Multiple reaction monitoring chromatograms of CL+30 (blue, 333.20/97.00; red, 333.20/201.00; green, 333.20/219.00; *m/z* in positive mode), CL+30 + 14 Da (blue, 347.00/223.00; red, 347.00/97.00; green, 347.00/205.00; *m/z* in positive mode) and CL+30 + 14 Da (blue, 345.00/301.00; red, 345.00/257.00; green, 345.00/113.00; *m/z* in negative mode) are shown.

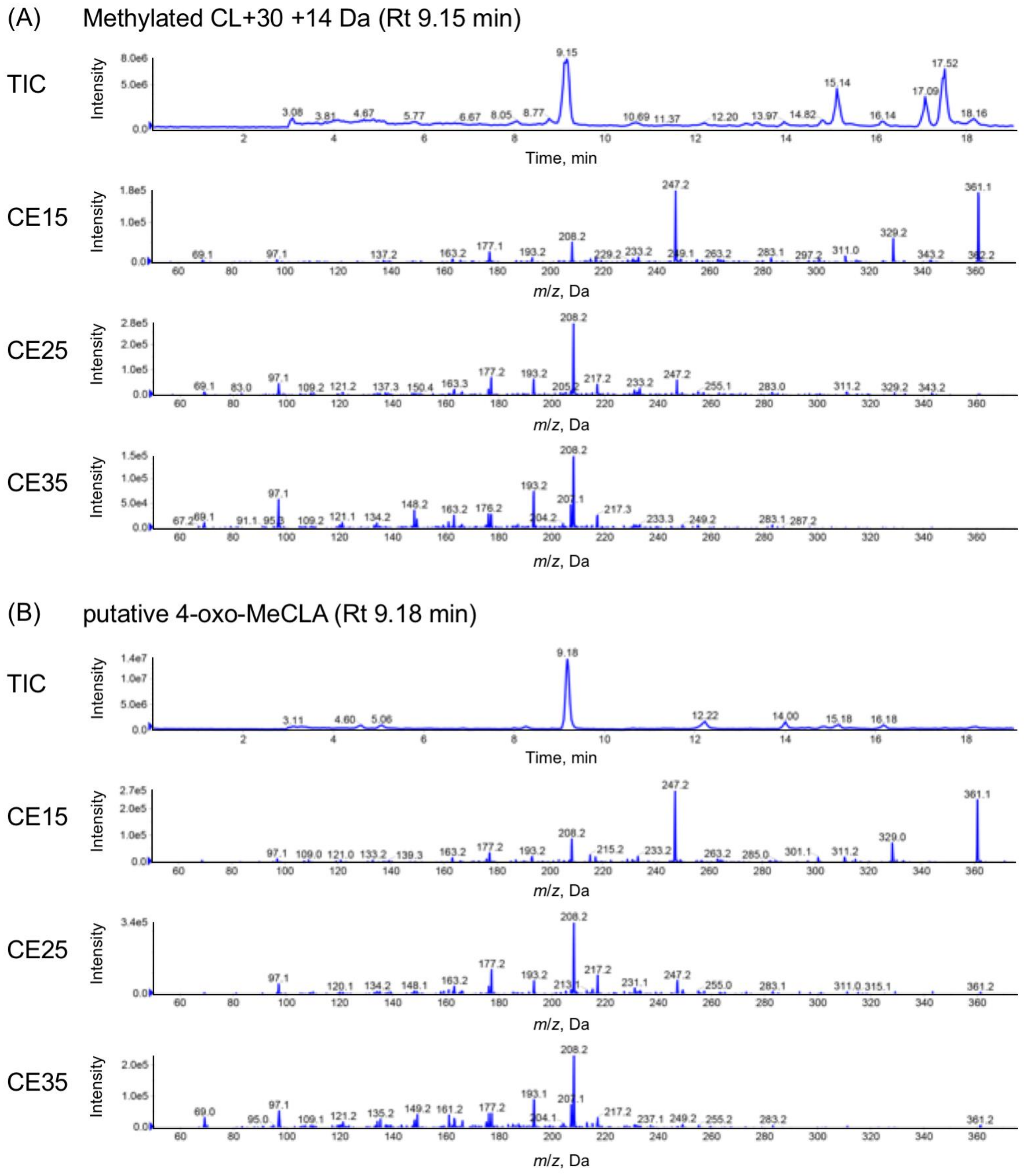


Fig. S7 Product ion spectra of (**A**) methylated CL+30 + 14 Da and (**B**) putative 4-oxo-MeCLA in root exudates of rice *d14* mutant. Total ion chromatogram (TIC) and product ion spectra derived from the precursor ion of *m*/*z* 361 $[M+H]^+$ in positive mode by Collision Energy (CE) 15, 25 and 35 V were shown.

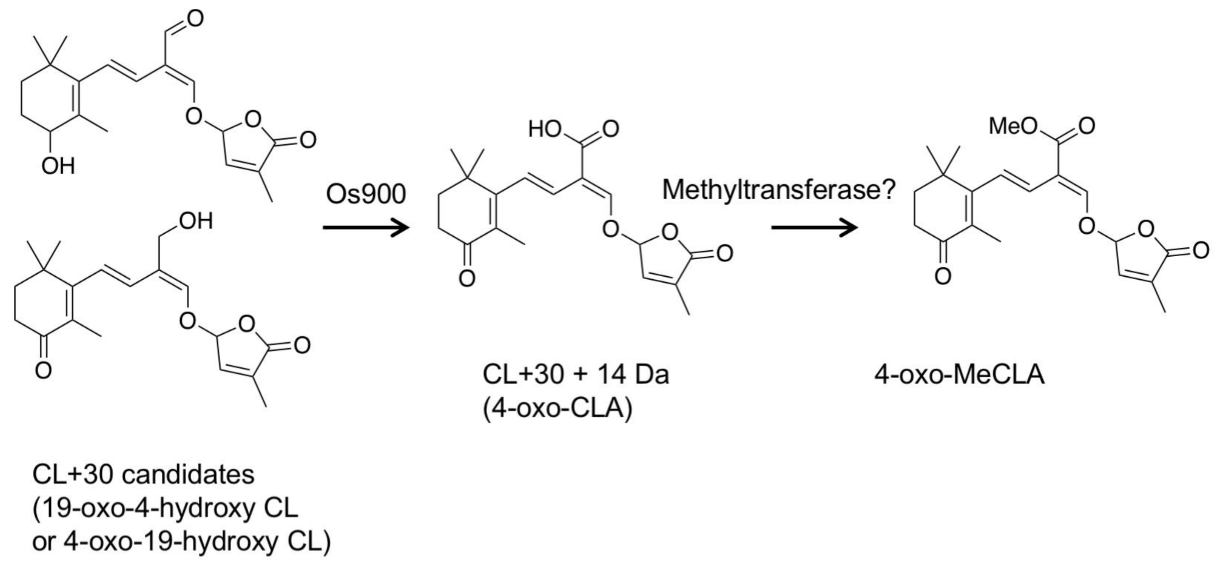


Fig. S8 Proposed 4-oxo-MeCLA biosynthesis. Oxidation at C4 and C19 position by Os900 gives 4-oxo-CLA. The conversion of 4-oxo-CLA to tentative 4-oxo-MeCLA is catalyzed by unknown methyltransferase

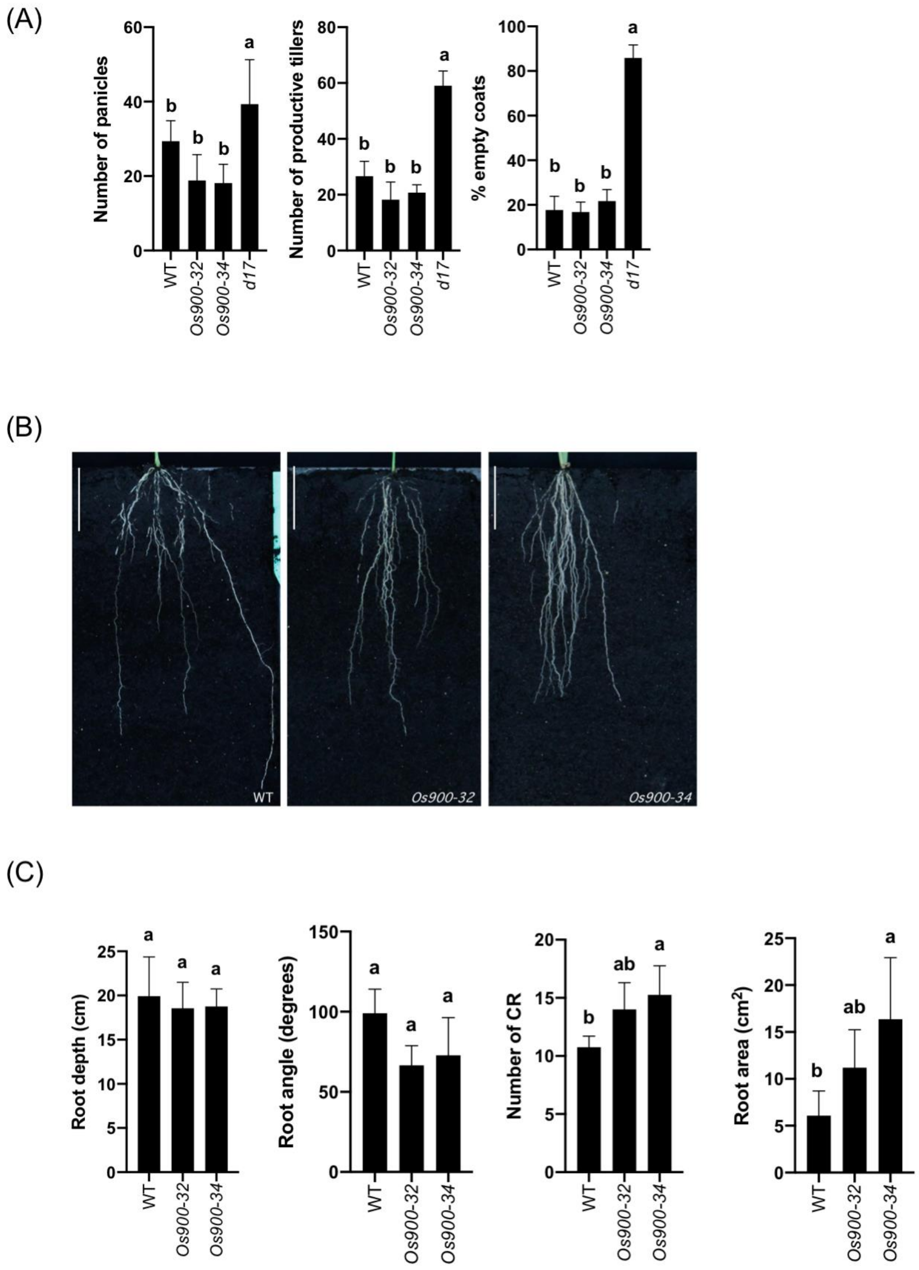


Fig. S9 Phenotypes of Os900-KO lines. (**A**) Shoot phenotypes of WT, Os900KO-lines and d17 mutant grown in soil. The data are represented as mean ± SD for a number of biological replicates (5≤n≤7 for WT, *Os900-32* and -*34*, n=3 for *d17*). (**B** and **C**) Root phenotypes of WT and Os900-KO lines. The data are represented as mean ± SD for a number of samples n=4. The statistical significance is determined by one-way ANOVA and Tukey's multiple comparison test.

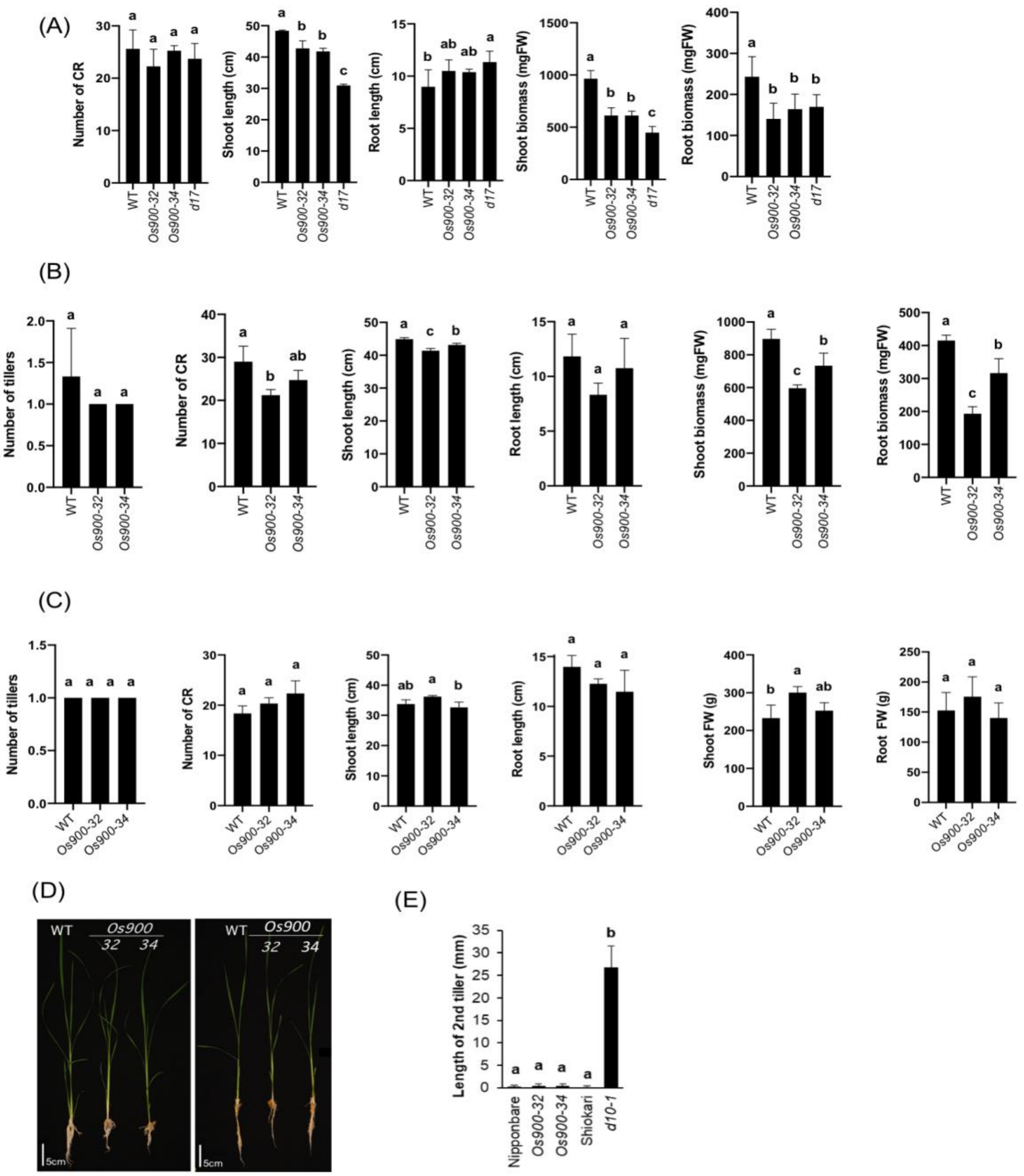

Fig. S10 Phenotypes of *Os900-KO* lines grown in the hydroponic culture under (**A**) +P, (**B**) –P, and (**C**) low Pi conditions. (**D**) Pictures of 3-week old seedlings grown in –Pi (left) and low Pi (right). (**E**) Second tiller of two-week-old rice SL biosynthetic mutants *Os900*-KO lines 32 and 34, and *d10-1*. The data are presented as mean ± SD for the number of biological replicates (A, n=5 for WT, n=8 for *Os900-32*, n=4 for *Os900-34*, and n=7 for *d17*; B and C, 3≤n≤5; E, n=10). The statistical significance is determined by one-way ANOVA and Tukey's multiple comparison test.

(A)

(B)

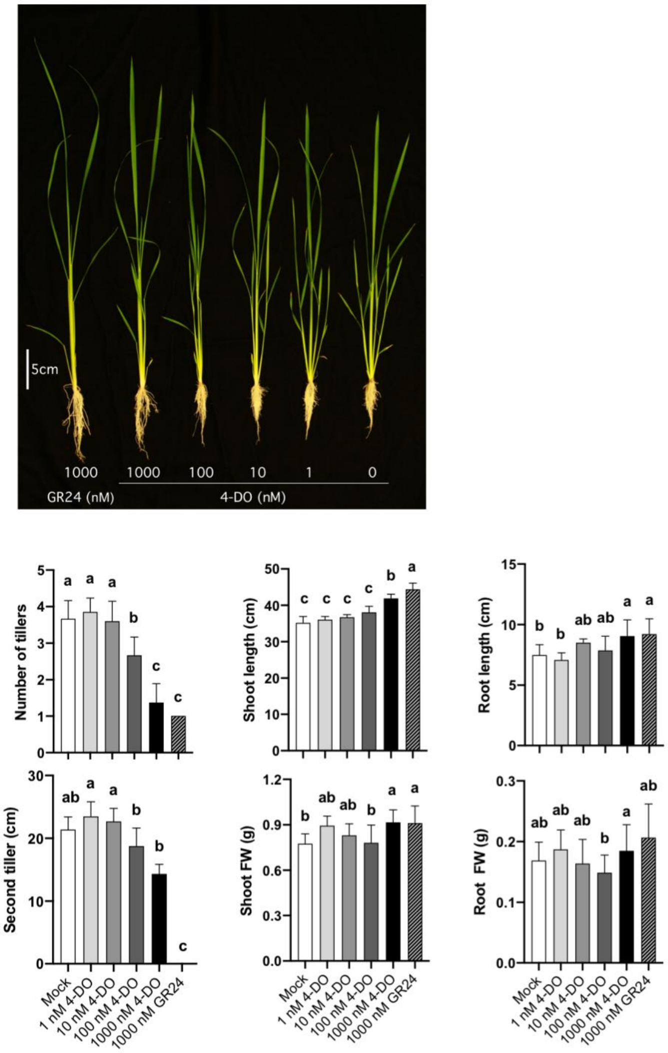


Fig. S11 Effect of 4-deoxyorobanchol (4DO) on the tillers, shoot and root growth of *d17* SL deficient mutant. The values are represented as the mean ± SD number of biological replicates (5≤n≤9). The statistical significance is determined by one-way ANOVA and Tukey's multiple comparison test.

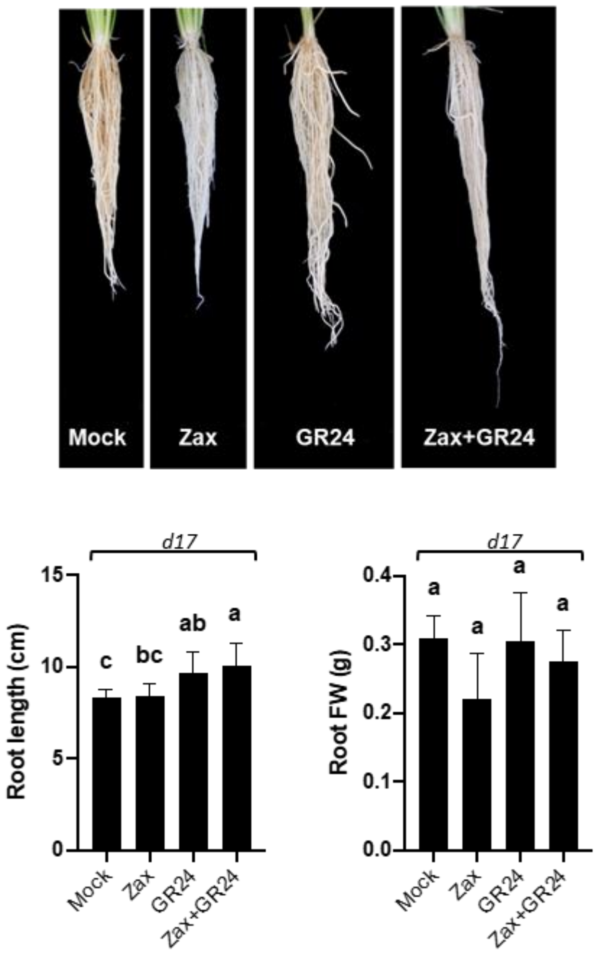


Fig. S12 Effects of 2.5 μM zaxinone with or without 1 μM *rac*-GR24 on the growth of *d17*. Data represent the mean ± SD for 6 biological replcates. Statistical analysis was performed using one-way analysis of variance (ANOVA) and Tukey's post hoc test. Different letters denote significant differences ($P < 0.05$).

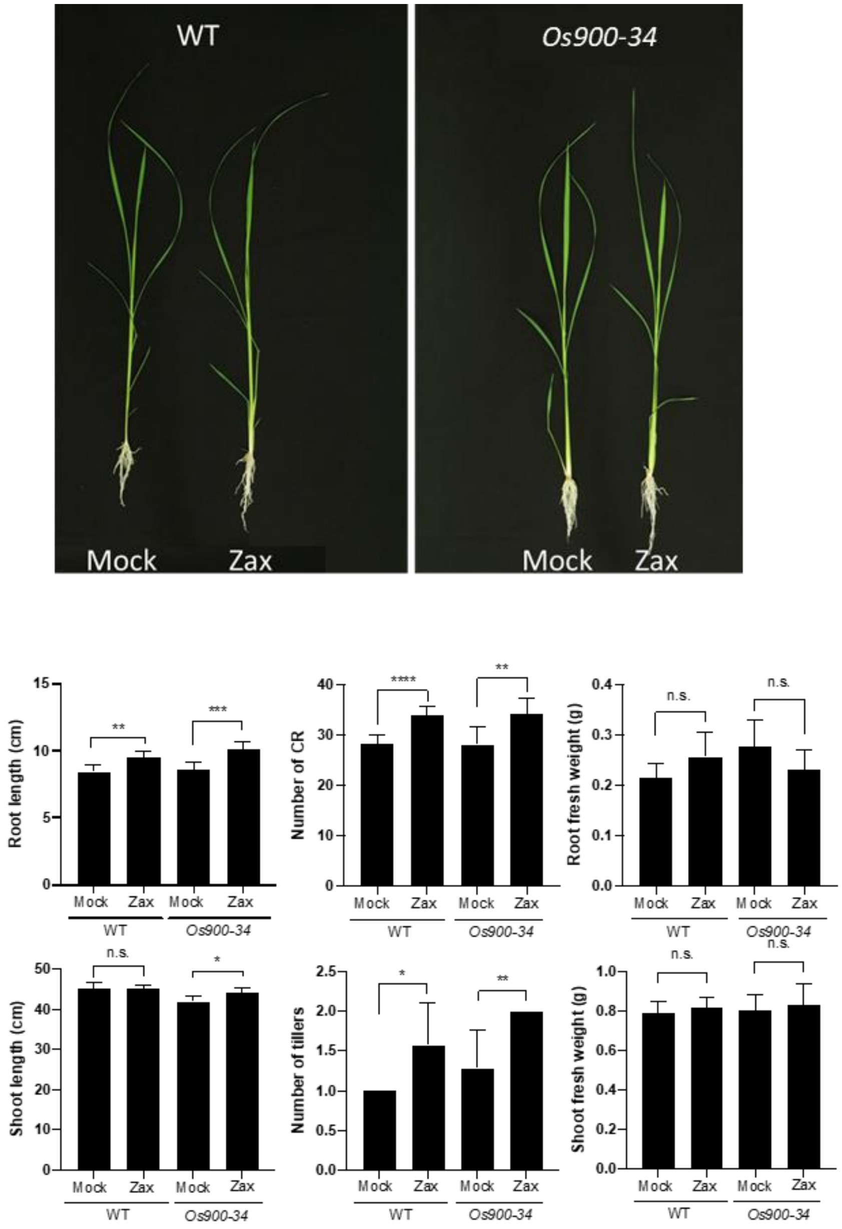


Fig. S13 Effects of 2.5 μM zaxinone (Zax) on the growth of *Os900*-KO compared to WT. Data represent the mean ± SD of 7 biological replicates. Statistical analysis was performed using one-way analysis of variance (ANOVA) and Tukey's post hoc test. (*P<0.05, **$P < 0.01$, ***$P < 0.001$, ****$P \leq 0.0001$, n.s. not significant)

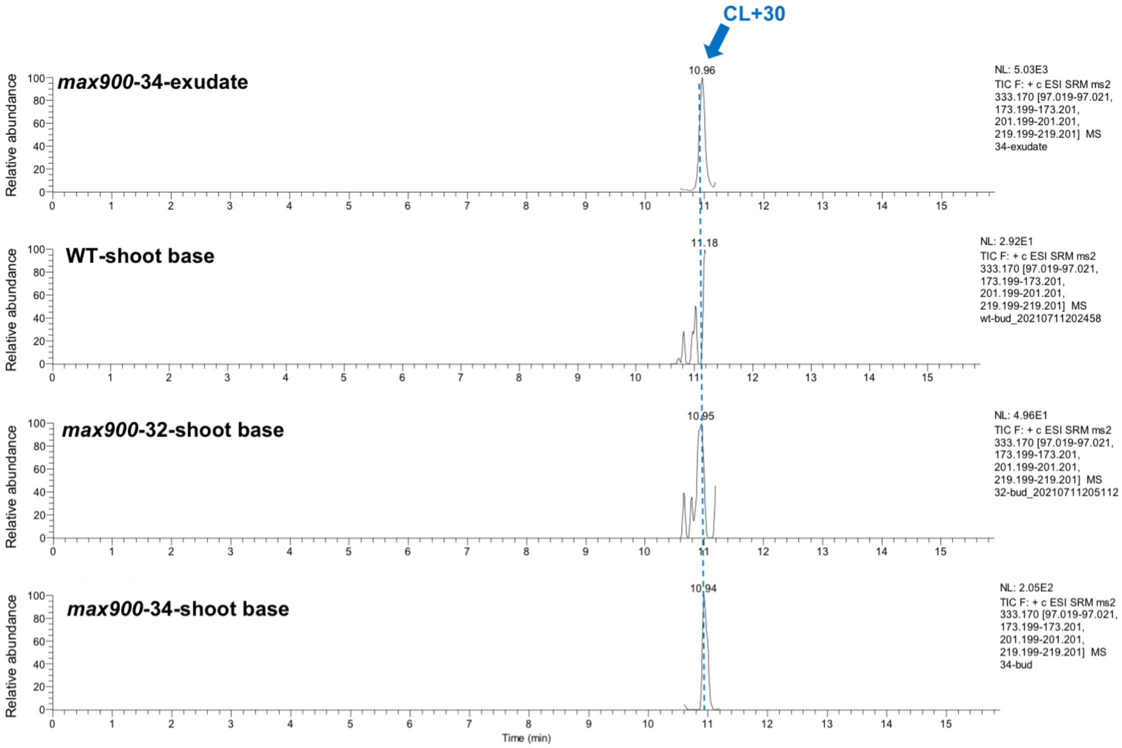


Fig. S14 Detection of CL+30 (4-oxo-hydroxyl-CL) in *Os900-34* root exudate and in the WT, *Os900-32* and *-34* shoot base (low Pi condition). (From up to down) Chromatograms of CL+30 detected in *Os900-34* root exudate, as well as in *Os900-32* and *-34* shoot base of plants grown in low Pi condition. No CL+30 detected in the shoot base of WT plants. The shoot base extraction samples were obtained from a pool of 12 shoot base of each genotype grown in low Pi condition. The blue arrows indicate the elution peaks of CL+30 with a retention time of 10.94 min.

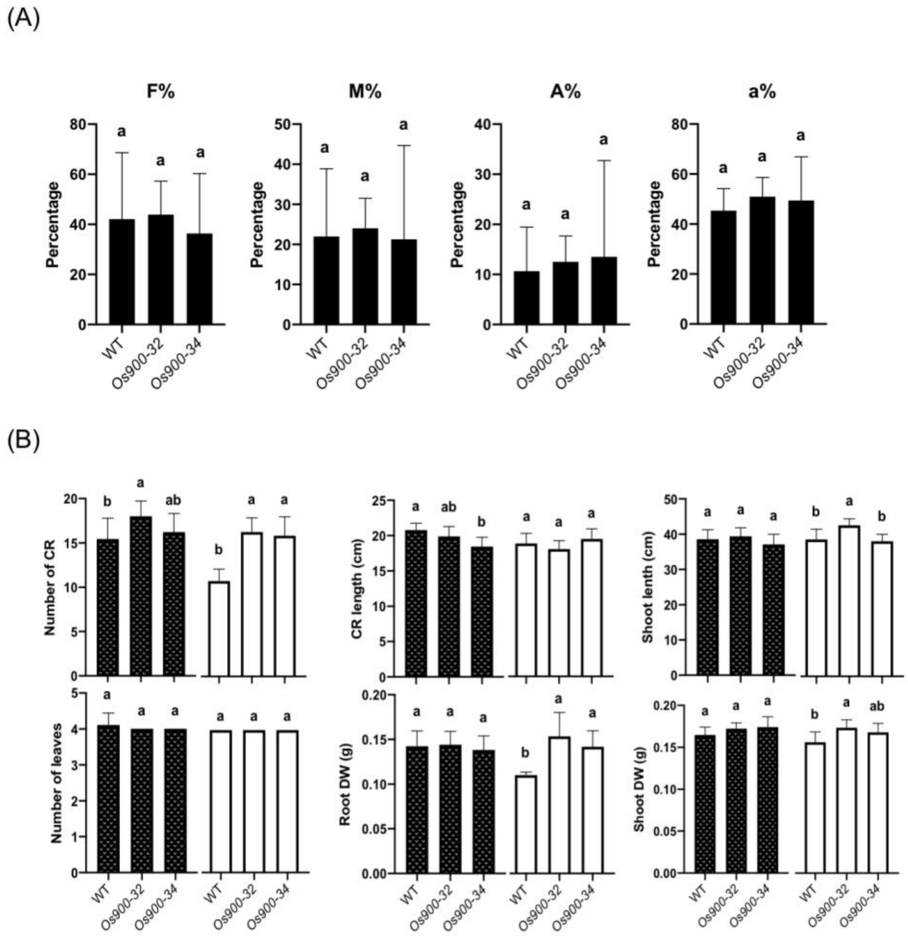


Fig. S15: Evaluation AM symbiosis colonization in *Osmax1* mutant lines. (**A**) Mycorrhizal colonization WT and *Os900*-KO lines by the AM fungus *Rhizophagus irregularis* at 35 dpi. Degree of colonization expressed as mycorrhizal frequency (F %), intensity (M %) and arbuscule abundance (A %); with the percentage of arbuscules within infected (a%) in the root system of WT and *Os900*-KO lines. Data are represented as the mean ± SD for 4 plants. (**B**) Plant phenotype comparison of plants with (plain bars) and without (white bars) mycorrhizal colonization by *R. irregularis* at 35 dpi for WT and *Os900*-KO lines. The collected traits were number of crown roots (CR), CR and shoot length, number of leaves, root and shoot dry weight (DW). The data are represented as the mean ± SD of 9 plants grown with AM fungi and 7 plants grown without. The statistical significance is determined by one-way ANOVA and Tukey's multiple comparison test.

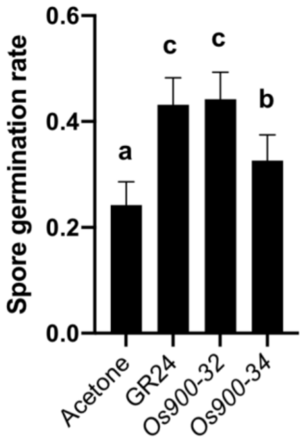


Fig. S16: Evaluation of *Osmax1-900* exudates on *Gigaspora margarita* spore germination. Acetone (Mock), GR24, and root exudates of *Os900-32* and *-34* were applied on *G. margarita* and the spore germination was reported. Data are the average of 96 biological replicates ± SD.

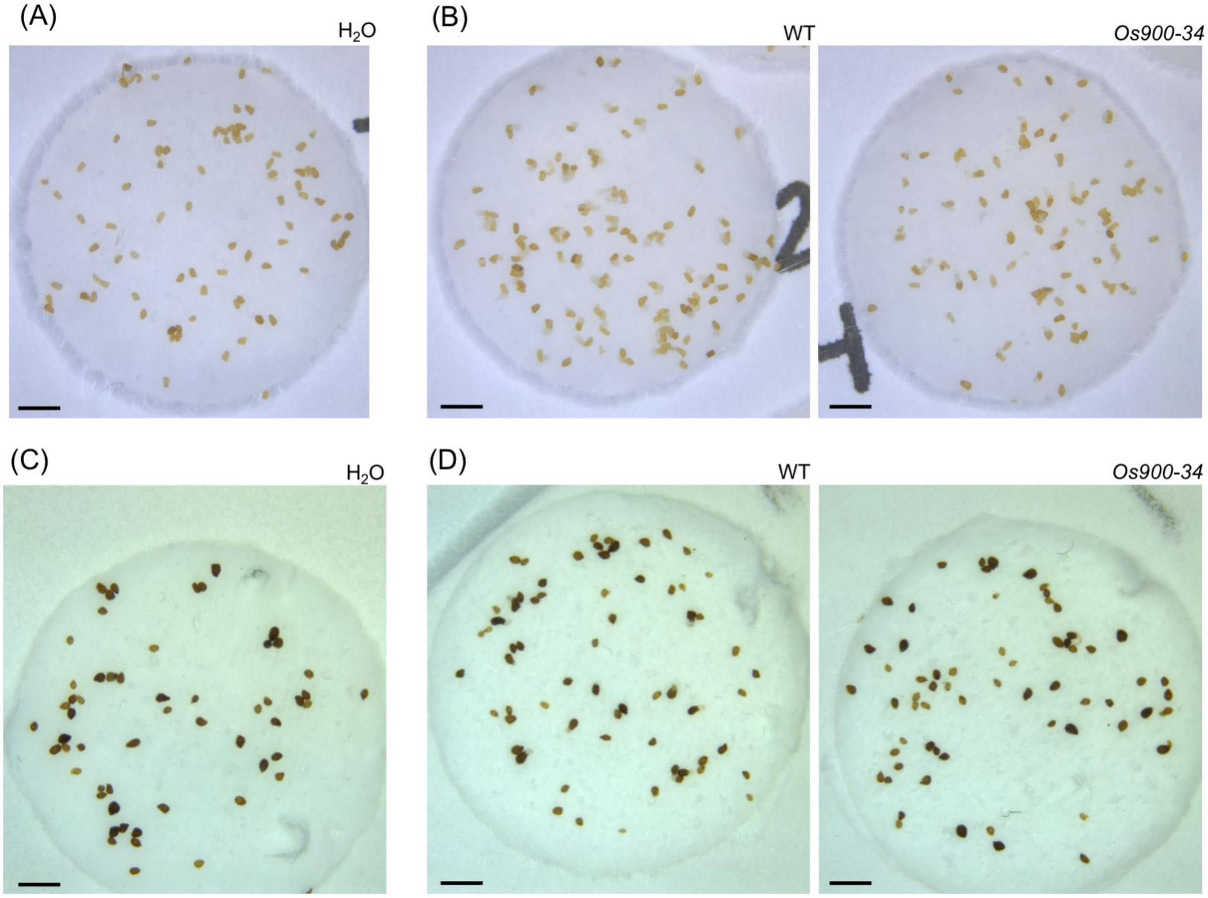


Fig. S17 Germination of the root parasitic weeds by treatment of root exudates. (**A**-**B**) on *Striga hermonthica* (**C**-**D**) on *Phelipanche ramose.*

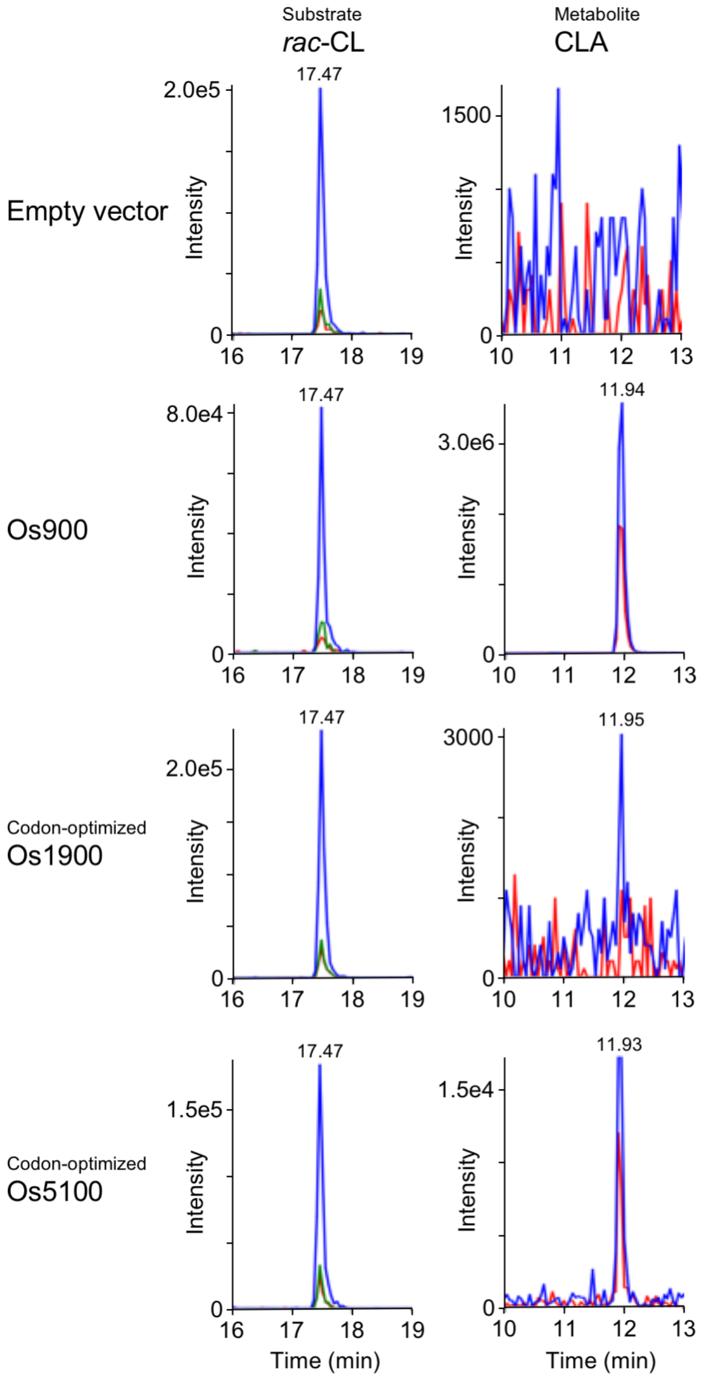


Fig. S18 Enzymatic conversion of CL to CLA by rice MAX1 homologs. Multiple reaction monitoring chromatograms of CL (blue, 303.00/97.00; red, 303.00/189.00; green, 303.00/207.00; *m/z* in positive mode) and CLA (blue, 331/113; red, 331/69; *m/z* in negative mode) are shown.

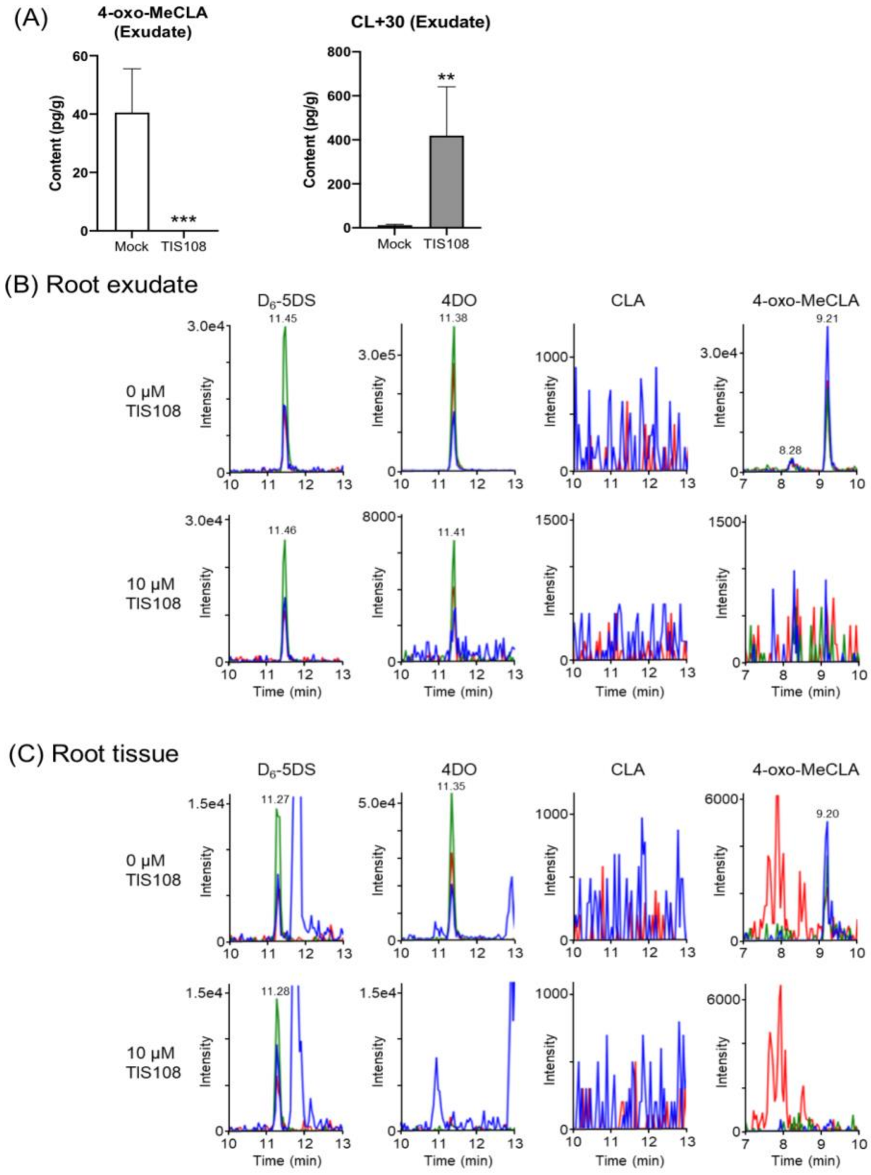


Fig. S19 Endogenous levels of SLs in wild-type rice root exudate (**A**-**B**) and tissues (**C**). Multiple reaction monitoring chromatograms of $D_6$-5DS (blue, 337/97; red, 337/222; green, 337/240; m/z in positive mode), 4DO (blue, 331/97; red, 331/216; green, 331/234; m/z in positive mode), CLA (blue, 331/113; red, 331/69; m/z in negative mode) and putative 4-oxo-MeCLA (blue, 361/208; red, 361/177; green, 361/97; m/z in positive mode) are shown. The data are presented as means ± SD of 5 biological replicates. Asterisk indicates significant difference without (Mock) and with 10 μM TIS108 treatment (TIS108) (***P*<0.01, ****P*<0.001, Student's t test).

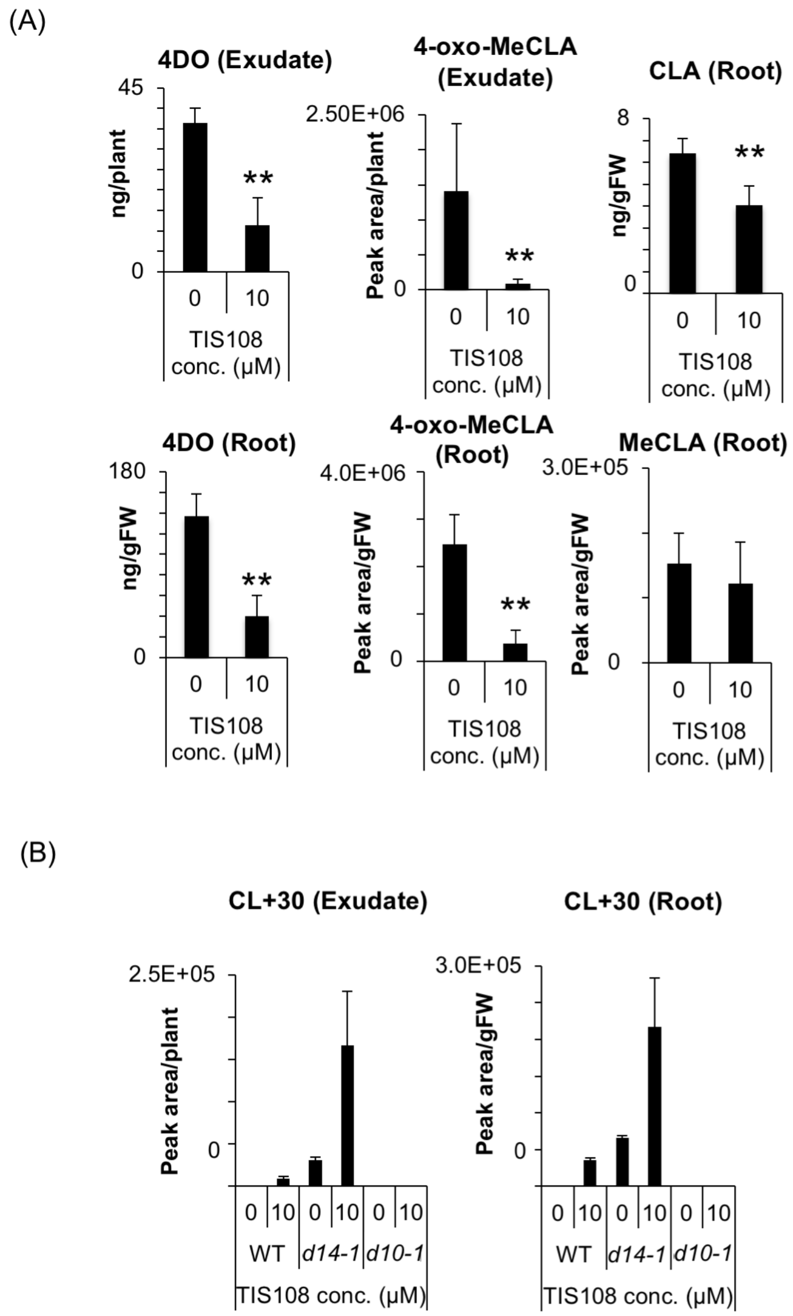


Fig. S20 Effects of TIS108 on endogenous SL level of *d14-1* rice. (**A**) Endogenous levels of SLs in roots and root exudates of the d14-1 mutant. The data are presented as means ± SD (n = 4). ** means statistically different from that of 0 μM TIS108 rice (t test, $P < 0.01$). (**B**) Endogenous levels of CL+30 in root exudates (left) and roots (right). The data are presented as means ± SD (n = 5).

(A)

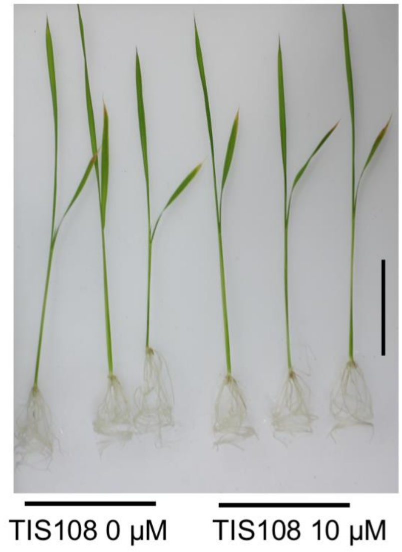


(B)

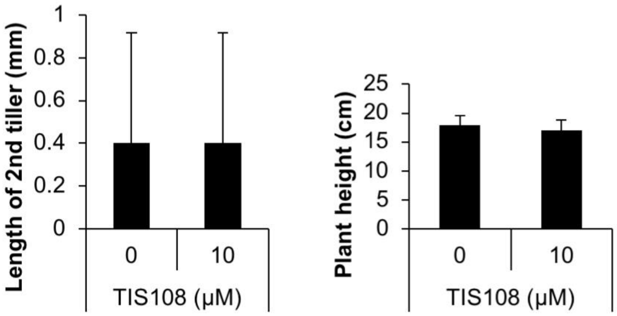


Fig. S21 Effect of TIS108 on two-week-old rice treated with or without 10 μM TIS108. (**A**) Phenotype on hydroponically grown seedlings. Scale bar = 5 cm. (**B**) Second tiller length (upper) and plant height (lower). The data are presented as means ± SD from 10 samples.

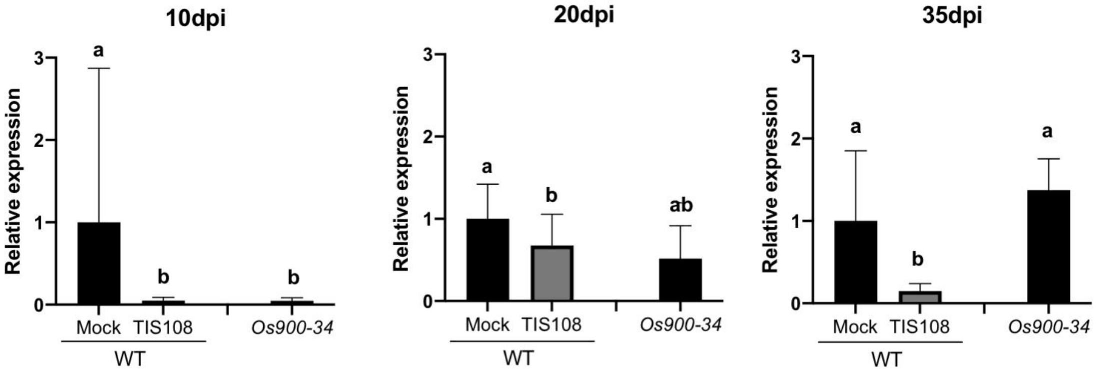


Fig. S22 Effect of TIS108 on mycorrhization. AM colonization was quantified by measuring the expression of *OsPT11* plant marker gene. The data are represented as the mean ± SD of number of samples n ($4 \leq n \leq 6$). The statistical significance is determined by Kruskal Wallis' test (non-parametric test).

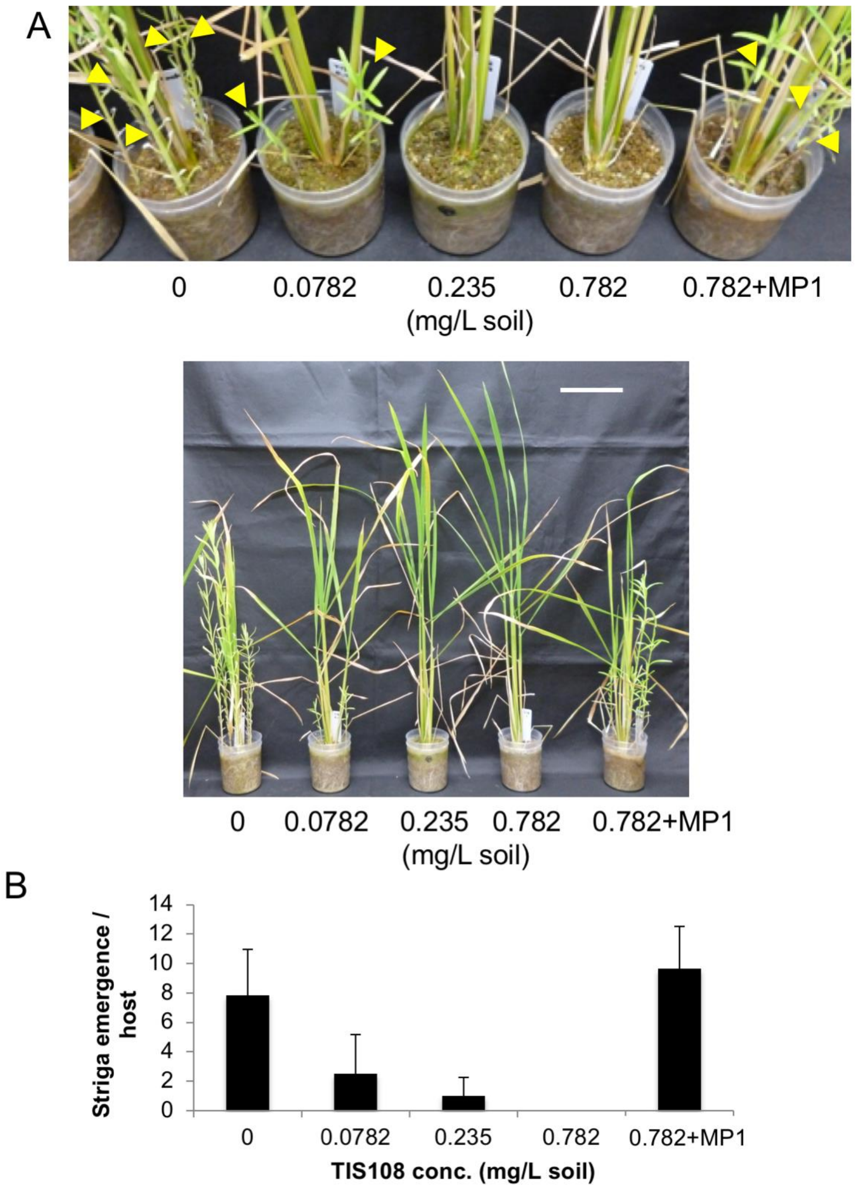


Fig. S23 *Striga* emergence test. Rice plants were co-cultivated with *Striga* for 7 weeks. The soil was weekly treated with acetone or TIS108. (**A**) Arrowheads indicate emerged *Striga*. Scale bar indicate 10 cm. (**B**) Number of emerged *Striga* after 7 weeks of co-cultivation. The data are means ± SD of 6 samples.

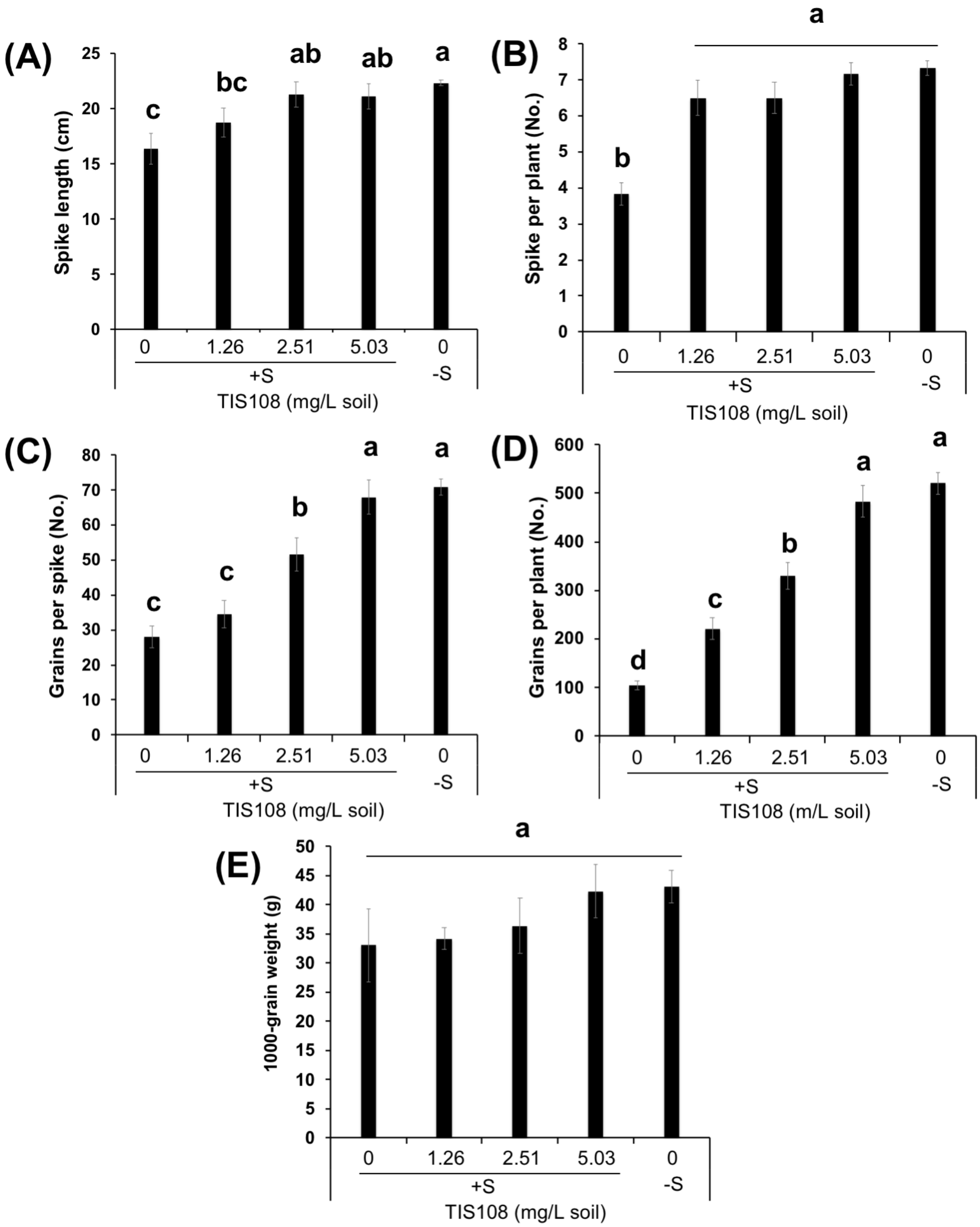


Fig. S24 Effects of TIS108 on rice growth and yield grown on the soil with (+S) or without (-S) *Striga* seeds. The soil was treated by TIS108 for three weeks at one-week interval. Spike length (**A**), number of spike (**B**), grains per spike (**C**), grains per plants (**D**) and 1000 grain weight. (**E**) were observed at the time of final harvesting. The data means ± SE of 6 samples. Means not sharing a letter in common differ significantly at $P_{0.05}$.

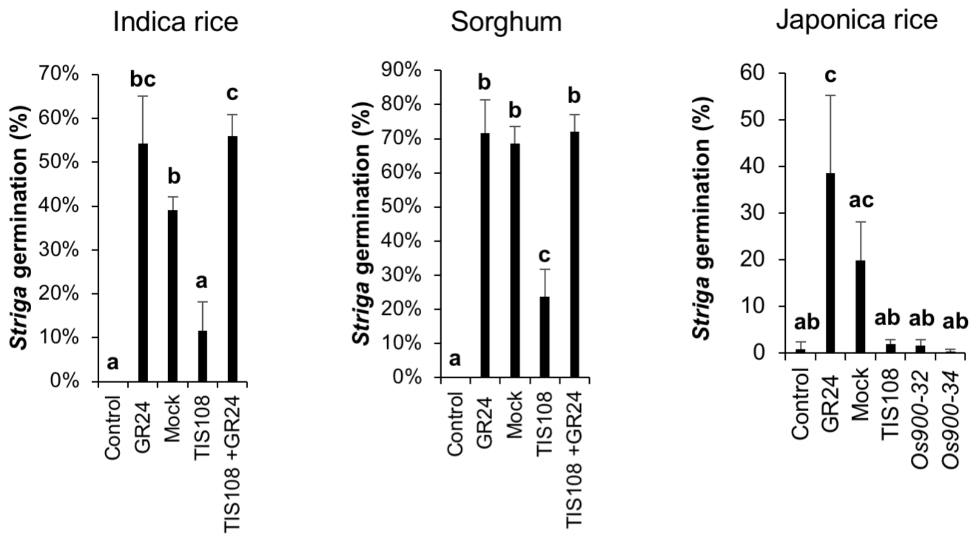


Fig. S25 *Striga* germination test in TIS108-treated Indica rice, sorghum, and Japonica rice. Control: water, GR24: 1 μM GR24, Mock: Root exudate of mock-treated plants, TIS108: Root exudate of 10 μM TIS108-treated plants, TIS108+GR24: Root exudate of 10 μM TIS108-treated plants and 1 μM GR24. The data are presented as means ± SD from 3-4 biological samples. Different letters indicate statistically significant differences at *P*0.05.

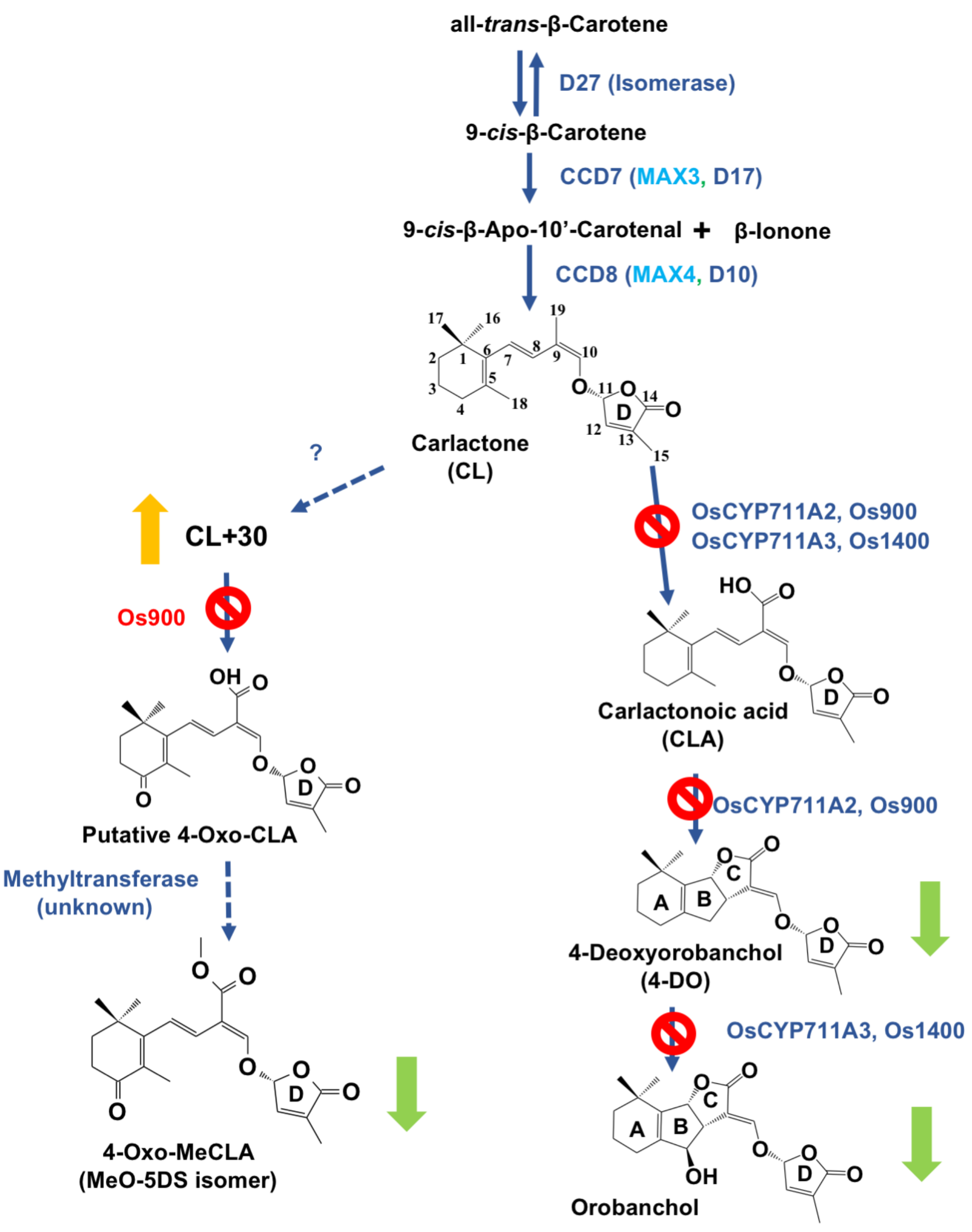


Fig. S26 TIS108, a MAX1-specific inhibitor of the SL biosynthesis. Abbreviations: D, Dwarf; CCD, Carotenoid Cleavage Dioxygenase; MAX, More Axillary Growth; Os900, OsMAX1-900; Os1400, OsMAX1-1400; CYP, Cytochrome P450; MeO-5DS, Methoxy-5-Deoxy-Strigol.

.

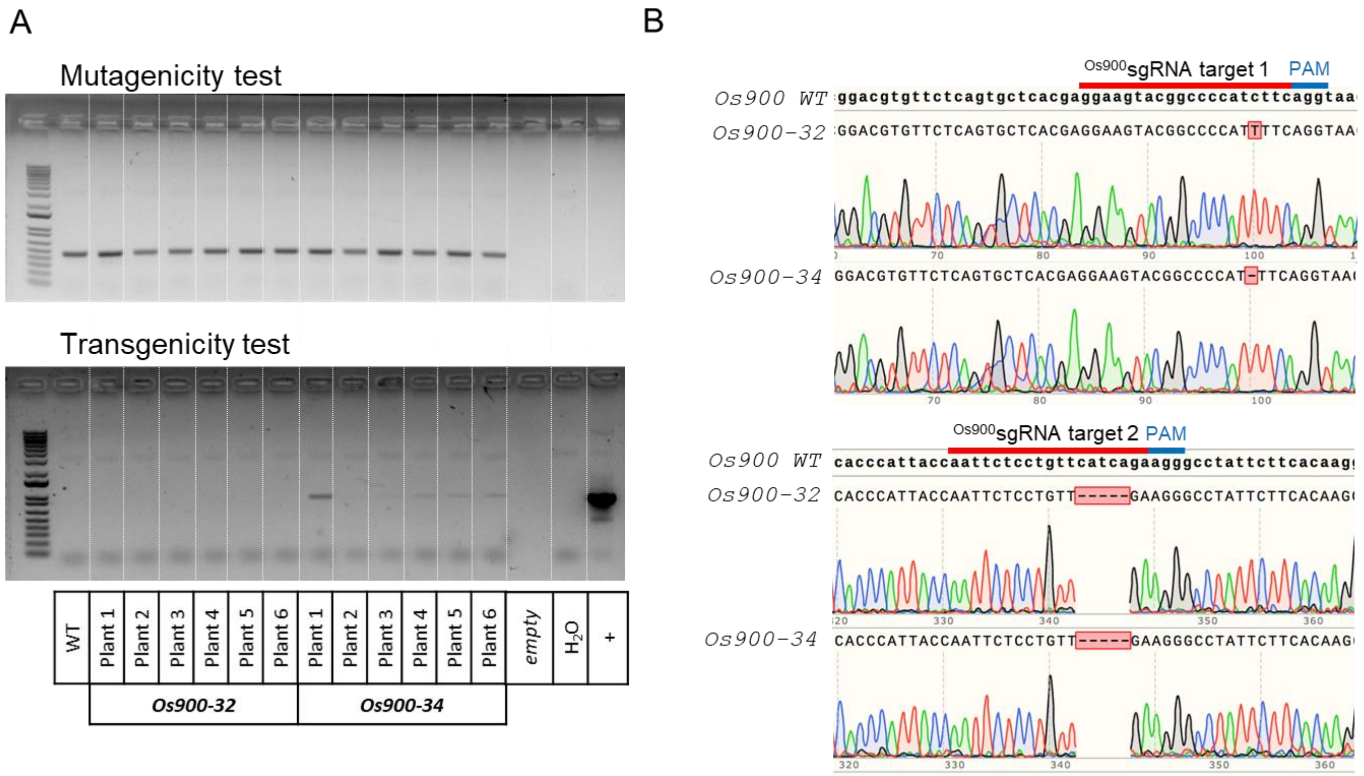


Fig. S27 Genotype details of Os*900*-KO lines. (**A**) Genomic DNA amplifications of the surrounding sgRNA target regions in wild-type (WT) and *Os900*-KO lines 32 and 34 (6 plants each) (up, mutagenicity test) and pRGEB32 region containing the two *Os900*sgRNAs sequences (down, transgenicity test). Water (H2O) was used as a negative control and pRGEB32 vector containing the two *Os900*sgRNAs sequences as a positive control (+). (**B**) Sequencing details of two representative plants of the homozygous *Os900*-KO lines showing the different mutations present in each line, aligned to the WT sequence for both *Os900* sgRNAs target sites.

### Table S1. Up-regulated genes of 10 μM TIS108-treated rice

| Gene Symbol | RAP ID | MSU ID | Description | Fold change |
|---|---|---|---|---|
| gene:Os04g0649700 | Os04g0649700 | LOC_Os04g55620.1 | receptor kinase, putative, expressed | 731.74 |
| gene:Os03g0784800 | Os03g0784800 | LOC_Os03g57130.1 | expressed protein | 592.28 |
| OsUMAMIT6 | Os02g0703900 | LOC_Os02g47500.1 | nodulin, putative, expressed | 464.19 |
| gene:Os05g0102800 | Os05g0102800 | LOC_Os05g01240.1 | AML1, putative, expressed | 80.88 |
| gene:Os02g0519950 | Os02g0519950 | None | | 69.14 |
| gene:Os10g0361000 | Os10g0361000 | LOC_Os10g21670.1 | dehydration stress-induced protein, putative, expressed | 64.29 |
| OsLAC8 | Os01g0850700 | LOC_Os01g63190.1 | laccase precursor protein, putative, expressed | 57.93 |
| gene:Os11g0137300 | Os11g0137300 | LOC_Os11g04220.1 | HEAT repeat family protein, putative, expressed | 39.23 |
| Cen8.t01003 | Os08g0314800 | LOC_Os08g22354.1 | polyadenylate-binding protein, putative, expressed | 37.24 |
| gene:Os03g0191100 | Os03g0191100 | LOC_Os03g09110.1 | mitochondrial carrier protein, putative, expressed | 21.51 |
| gene:Os01g0237750 | Os01g0237750 | LOC_Os01g13610.1 | isoflavone reductase homolog IRL, putative, expressed | 20.49 |
| OsPDR9 | Os01g0609300 | LOC_Os01g42380.1 | pleiotropic drug resistance protein, putative, expressed | 19.60 |
| gene:Os01g0760701 | Os01g0760701 | None | | 19.54 |
| OsPR10a | Os12g0555500 | LOC_Os12g36880.1 | pathogenesis-related Bet v I family protein, putative, expressed | 18.62 |
| OsABCG35 | Os01g0609200 | LOC_Os01g42370.1 | pleiotropic drug resistance protein, putative, expressed | 18.49 |
| gene:Os01g0237500 | Os01g0237500 | LOC_Os01g13610.1 | isoflavone reductase homolog IRL, putative, expressed | 17.51 |
| SSIVA | Os01g0720600 | LOC_Os01g52250.1 | starch synthase, putative, expressed | 17.36 |
| OsPDR9 | Os01g0609300 | LOC_Os01g42380.1 | pleiotropic drug resistance protein, putative, expressed | 16.99 |
| gene:Os01g0937012 | Os01g0937012 | None | | 15.20 |
| OsAMT2_2 | Os01g0831900 | LOC_Os01g61550.1 | ammonium transporter protein, putative, expressed | 15.14 |
| RAB21_2 | Os11g0454000 | LOC_Os11g26760.1 | dehydrin, putative, expressed | 13.70 |
| OsLEA25 | Os11g0451700 | LOC_Os11g26570.1 | dehydrin, putative, expressed | 13.67 |
| OsCYP709C9 | Os07g0418500 | LOC_Os07g23570.1 | cytochrome P450 72A1, putative, expressed | 13.66 |
| RAB21_1 | Os11g0453900 | LOC_Os11g26750.1 | dehydrin, putative, expressed | 12.25 |
| gene:Os01g0609501 | Os01g0609501 | None | | 12.18 |
| RAB16B | Os11g0454200 | LOC_Os11g26780.1 | dehydrin, putative, expressed | 11.14 |
| gene:Os11g0311300 | Os11g0311300 | LOC_Os11g20689.1 | exosome complex exonuclease, putative, expressed | 9.47 |
| gene:Os05g0513900 | Os05g0513900 | LOC_Os05g43830.1 | hydrolase, alpha/beta fold family domain containing protein, expressed | 9.36 |
| gene:Os05g0410601 | Os05g0410601 | None | | 9.10 |
| OsEnS-53 | Os03g0723400 | LOC_Os03g51350.1 | expressed protein | 8.84 |
| Oschib1 | Os10g0416500 | LOC_Os10g28080.1 | glycosyl hydrolase, putative, expressed | 8.62 |
| gene:Os05g0151200 | Os05g0151200 | LOC_Os05g05930.1 | peripheral-type benzodiazepine receptor, putative, expressed | 8.29 |
| OsLEA16 | Os03g0168100 | LOC_Os03g07180.1 | embryonic protein DC-8, putative, expressed | 6.91 |
| Rab16A | Os11g0454300 | LOC_Os11g26790.1 | dehydrin, putative, expressed | 6.77 |
| gene:Os06g0691700 | Os06g0691700 | None | | 6.76 |
| OsMaT-2 | Os02g0483500 | LOC_Os02g28170.1 | transferase family protein, putative, expressed | 6.52 |
| gene:Os03g0429800 | Os03g0429800 | LOC_Os03g31550.1 | aldehyde oxidase, putative, expressed | 6.49 |
| gene:Os03g0792850 | Os03g0792850 | None | | 6.35 |
| RSOsPR10 | Os12g0555000 | LOC_Os12g36830.1 | pathogenesis-related Bet v I family protein, putative, expressed | 6.12 |
| OsKMD3 | Os02g0563000 | LOC_Os02g35530.1 | OsFBK8 - F-box domain and kelch repeat containing protein, expressed | 5.92 |
| gene:Os07g0585900 | Os07g0585900 | LOC_Os07g39720.1 | expressed protein | 5.89 |
| OsTHI9 | Os06g0514800 | LOC_Os06g32240.1 | THION9 - Plant thionin family protein precursor, expressed | 5.66 |
| gene:Os01g0639600 | Os01g0639600 | LOC_Os01g45250.1 | DUF1645 domain containing protein, putative, expressed | 5.48 |
| OsGH3-7 | Os06g0499500 | LOC_Os06g30440.1 | OsGH3.7 - Probable indole-3-acetic acid-amido synthetase, expressed | 5.39 |
| OsCCR10 | Os02g0811800 | LOC_Os02g56700.1 | dehydrogenase, putative, expressed | 5.31 |
| gene:Os04g0653700 | Os04g0653700 | LOC_Os04g55980.1 | glycine-rich RNA-binding, abscisic acid-inducible protein, putative, expressed | 5.22 |
| OsSub31 | Os03g0761500 | LOC_Os03g55350.1 | OsSub31 - Putative Subtilisin homologue, expressed | 5.14 |
| gene:Os05g0526700 | Os05g0526700 | LOC_Os05g45070.1 | harpin-induced protein 1 domain containing protein, expressed | 5.14 |
| gene:Os02g0207800 | Os02g0207800 | None | | 5.07 |
| AWPM-19 | Os05g0381400 | LOC_Os05g31670.1 | AWPM-19-like membrane family protein, putative, expressed | 5.01 |
| Cht6 | Os02g0605900 | LOC_Os02g39330.1 | CHIT1 - Chitinase family protein precursor, expressed | 4.97 |
| ABCG43 | Os07g0522500 | LOC_Os07g33780.1 | pleiotropic drug resistance protein 5, putative, expressed | 4.85 |
| SPL7 | Os05g0530400 | LOC_Os05g45410.1 | HSF-type DNA-binding domain containing protein, expressed | 4.83 |
| gene:Os06g0129900 | Os06g0129900 | LOC_Os06g03930.1 | cytochrome P450 86A1, putative, expressed | 4.79 |
| RPR10b | Os12g0555200 | LOC_Os12g36850.1 | pathogenesis-related Bet v I family protein, putative, expressed | 4.49 |
| gene:Os01g0838600 | Os01g0838600 | LOC_Os01g62130.1 | ZOS1-14 - C2H2 zinc finger protein, expressed | 4.42 |
| gene:Os06g0239200 | Os06g0239200 | LOC_Os06g13190.1 | expressed protein | 4.39 |
| gene:Os01g0644200 | Os01g0644200 | None | | 4.34 |
| gene:Os06g0129900 | Os06g0129900 | LOC_Os06g03930.1 | cytochrome P450 86A1, putative, expressed | 4.31 |
| C10728 | Os01g0660200 | LOC_Os01g47070.1 | glycosyl hydrolase, putative, expressed | 4.26 |
| gene:Os03g0326200 | Os03g0326200 | LOC_Os03g20970.1 | phospholipid-transporting ATPase 1, putative, expressed | 4.15 |
| OsMAS | Os04g0179200 | LOC_Os04g10010.1 | sex determination protein tasselseed-2, putative, expressed | 4.11 |
| gene:Os10g0535800 | Os10g0535800 | LOC_Os10g39100.1 | uncharacterized Cys-rich domain containing protein, putative, expressed | 4.10 |
| gene:Os02g0102400 | Os02g0102400 | LOC_Os02g01230.1 | ribosomal protein, putative, expressed | 3.95 |
| gene:Os07g0568000 | Os07g0568000 | LOC_Os07g38060.1 | expressed protein | 3.93 |
| OsMSRA2.2 | Os04g0482100 | LOC_Os04g40620.1 | peptide methionine sulfoxide reductase, putative, expressed | 3.92 |
| gene:Os07g0635150 | Os07g0635150 | None | | 3.86 |
| gene:Os03g0351700 | Os03g0351700 | LOC_Os03g22820.1 | expressed protein | 3.81 |

#### Table S1. Continued

| | | | | |
|---|---|---|---|---|
| gene:Os08g0173600 | Os08g0173600 | LOC_Os08g07690.1 | expressed protein | 3.68 |
| gene:Os01g0934300 | Os01g0934300 | LOC_Os01g70810.1 | homeobox domain containing protein, expressed | 3.68 |
| gene:Os11g0514500 | Os11g0514500 | LOC_Os11g31540.1 | BRASSINOSTEROID INSENSITIVE 1-associated receptor kinase 1 precursor, putative, expressed | 3.65 |
| gene:Os05g0444200 | Os05g0444200 | LOC_Os05g37190.1 | ZOS5-08 - C2H2 zinc finger protein, expressed | 3.57 |
| gene:Os04g0173800 | Os04g0173800 | LOC_Os04g09390.1 | HEV3 - Hevein family protein precursor, expressed | 3.50 |
| OsSCP64 | Os11g0643400 | LOC_Os11g42390.1 | OsSCP64 - Putative Serine Carboxypeptidase homologue, expressed | 3.49 |
| gene:Os02g0209400 | Os02g0209400 | LOC_Os02g11870.1 | expressed protein | 3.46 |
| gene:Os01g0660000 | Os01g0660000 | None | | 3.46 |
| gene:Os05g0546400 | Os05g0546400 | LOC_Os05g46840.1 | proline-rich protein, putative, expressed | 3.44 |
| gene:Os01g0341750 | Os01g0341750 | None | | 3.43 |
| gene:Os05g0112800 | Os05g0112800 | LOC_Os05g02200.1 | cysteine-rich repeat secretory protein 55 precursor, putative, expressed | 3.42 |
| gene:Os05g0522600 | Os05g0522600 | LOC_Os05g44770.1 | receptor-like protein kinase 5 precursor, putative, expressed | 3.38 |
| gene:Os07g0538000 | Os07g0538000 | LOC_Os07g35350.1 | glucan endo-1,3-beta-glucosidase precursor, putative, expressed | 3.37 |
| gene:Os07g0125201 | Os07g0125201 | LOC_Os07g03319.1 | SCP-like extracellular protein, expressed | 3.27 |
| OsPR1#072 | Os07g0127500 | LOC_Os07g03580.1 | SCP-like extracellular protein, expressed | 3.27 |
| gene:Os07g0635200 | Os07g0635200 | LOC_Os07g44110.1 | cytochrome P450 72A1, putative, expressed | 3.26 |
| gene:Os07g0126500 | Os07g0126500 | LOC_Os07g03499.1 | SCP-like extracellular protein, expressed | 3.24 |
| OsPYL/RCAR2 | Os02g0226801 | LOC_Os02g13330.1 | bet v I allergen family protein, putative, expressed | 3.19 |
| OsABCG48 | Os11g0587600 | LOC_Os11g37700.1 | pleiotropic drug resistance protein, putative, expressed | 3.15 |
| gene:Os12g0629700 | Os12g0629700 | None | | 3.11 |
| brd1 | Os03g0602300 | LOC_Os03g40540.1 | cytochrome P450, putative, expressed | 3.09 |
| gene:Os02g0209300 | Os02g0209300 | LOC_Os02g11859.1 | expressed protein | 3.07 |
| gene:Os05g0403300 | Os05g0403300 | LOC_Os05g33430.1 | xyloglucanase inhibitor, putative, expressed | 3.01 |
| RASI | Os04g0526600 | LOC_Os04g44470.1 | KUN1 - Kunitz-type trypsin inhibitor precursor, expressed | 3.01 |
| gene:Os05g0130100 | Os05g0130100 | LOC_Os05g03920.1 | TKL_IRAK_DUF26-lf.3 - DUF26 kinases have homology to DUF26 containing loci, expressed | 3.01 |
| OsABCG44 | Os08g0384500 | LOC_Os08g29570.1 | pleiotropic drug resistance protein 3, putative, expressed | 2.98 |
| gene:Os04g0634000 | Os04g0634000 | LOC_Os04g54140.1 | receptor-like kinase, putative, expressed | 2.94 |
| OsGELP11 | Os01g0216900 | LOC_Os01g11790.1 | GDSL-like lipase/acylhydrolase, putative, expressed | 2.94 |
| OsGELP57 | Os04g0561800 | LOC_Os04g47390.1 | GDSL-like lipase/acylhydrolase, putative, expressed | 2.93 |
| OsPR5_4 | Os12g0628600 | LOC_Os12g43380.1 | thaumatin, putative, expressed | 2.91 |
| OsLEA3-1 | Os05g0542500 | LOC_Os05g46480.1 | late embryogenesis abundant protein, group 3, putative, expressed | 2.82 |
| gene:Os02g0740600 | Os02g0740600 | LOC_Os02g50710.1 | expressed protein | 2.77 |
| RF2b | Os03g0336200 | LOC_Os03g21800.1 | bZIP transcription factor family protein, putative, expressed | 2.76 |
| gene:Os05g0576600 | Os05g0576600 | LOC_Os05g50100.1 | expressed protein | 2.75 |
| OsKOD1 | Os04g0507950 | None | | 2.75 |
| prx92 | Os06g0695300 | LOC_Os06g48010.1 | peroxidase precursor, putative, expressed | 2.69 |
| ADC2 | Os04g0107600 | LOC_Os04g01690.1 | pyridoxal-dependent decarboxylase protein, putative, expressed | 2.66 |
| gene:Os07g0126100 | Os07g0126100 | LOC_Os07g03409.1 | SCP-like extracellular protein, expressed | 2.64 |
| gene:Os06g0715500 | Os06g0715500 | LOC_Os06g50154.1 | translocon-associated protein subunit alpha precursor, putative, expressed | 2.64 |
| OsRLCK255 | Os08g0457400 | LOC_Os08g35600.1 | tyrosine protein kinase domain containing protein, putative, expressed | 2.63 |
| PDIL1_2 | Os02g0554900 | LOC_Os02g34940.1 | OsPDIL1-3 protein disulfide isomerase PDIL1-3, expressed | 2.61 |
| OsNADPH1 | Os04g0497000 | LOC_Os04g41960.1 | NADP-dependent oxidoreductase, putative, expressed | 2.61 |
| gene:Os02g0831300 | Os02g0831300 | LOC_Os02g58460.1 | beta-catenin-like protein 1, putative, expressed | 2.58 |
| OsLOX10 | Os11g0575600 | LOC_Os11g36719.1 | lipoxygenase, putative, expressed | 2.52 |
| OsASNase2 | Os04g0650700 | LOC_Os04g55710.1 | transposon protein, putative, unclassified, expressed | 2.51 |
| OsPR4d | Os11g0591800 | LOC_Os11g37940.1 | WIP2 - Wound-induced protein precursor, expressed | 2.51 |
| gene:Os02g0205500 | Os02g0205500 | LOC_Os02g11070.1 | 3-ketoacyl-CoA synthase, putative, expressed | 2.50 |
| gene:Os11g0246166 | Os11g0246166 | None | | 2.48 |
| HB1 | Os03g0233900 | LOC_Os03g13140.1 | non-symbiotic hemoglobin 2, putative, expressed | 2.47 |
| OsPLDalpha4 | Os06g0604200 | LOC_Os06g40170.1 | phospholipase D, putative, expressed | 2.46 |
| OsPGL32 | Os01g0623600 | LOC_Os01g43490.1 | polygalacturonase, putative, expressed | 2.44 |
| gene:Os04g0326100 | Os04g0326100 | LOC_Os04g25970.1 | cytokinin-O-glucosyltransferase 2, putative, expressed | 2.42 |
| gene:Os08g0158200 | Os08g0158200 | LOC_Os08g06170.1 | berberine and berberine like domain containing protein, expressed | 2.42 |
| gene:Os09g0564400 | Os09g0564400 | LOC_Os09g39090.1 | vignain precursor, putative, expressed | 2.34 |
| gene:Os06g0211600 | Os06g0211600 | LOC_Os06g10910.1 | xyloglucan fucosyltransferase, putative, expressed | 2.34 |
| OsDjC29 | Os03g0323600 | LOC_Os03g20730.1 | chaperone protein dnaJ, putative, expressed | 2.34 |
| OsPR5_2 | Os03g0661600 | LOC_Os03g45960.1 | thaumatin, putative, expressed | 2.34 |
| OsCDAP1 | Os07g0162400 | LOC_Os07g06830.1 | gibberellin receptor GID1L2, putative, expressed | 2.32 |
| NAC122 | Os11g0126900 | LOC_Os11g03300.1 | NAC domain transcription factor, putative, expressed | 2.30 |
| gene:Os03g0772600 | Os03g0772600 | LOC_Os03g56160.1 | lectin-like receptor kinase 7, putative, expressed | 2.30 |
| OsFKBP57 | Os01g0562400 | LOC_Os01g38180.1 | peptidyl-prolyl isomerase, putative, expressed | 2.27 |
| OsKMD4 | Os11g0246200 | LOC_Os11g14140.1 | OsFBK25 - F-box domain and kelch repeat containing protein, expressed | 2.26 |
| gene:Os03g0757000 | Os03g0757000 | LOC_Os03g55010.1 | UDP-glucoronosyl and UDP-glucosyl transferase domain containing protein, expressed | 2.26 |
| gene:Os03g0388600 | Os03g0388600 | LOC_Os03g27090.1 | MYB family transcription factor, putative, expressed | 2.25 |
| gene:Os10g0439100 | Os10g0439100 | LOC_Os10g30330.1 | expansin precursor, putative, expressed | 2.25 |
| gene:Os08g0267300 | Os08g0267300 | LOC_Os08g16660.1 | aspartic proteinase nepenthesin precursor, putative, expressed | 2.24 |
| gene:Os03g0726800 | Os03g0726800 | LOC_Os03g51670.1 | serine esterase family protein, putative, expressed | 2.23 |
| DEFL12 | Os04g0522100 | LOC_Os04g44130.1 | DEF12 - Defensin and Defensin-like DEFL family, expressed | 2.22 |
| OsGSTU41 | Os01g0950000 | LOC_Os01g72160.1 | glutathione S-transferase, putative, expressed | 2.22 |
| OsRMC | Os04g0659300 | LOC_Os04g56430.1 | cysteine-rich receptor-like protein kinase, putative, expressed | 2.20 |
| gene:Os08g0111200 | Os08g0111200 | LOC_Os08g01940.1 | non-lysosomal glucosylceramidase, putative, expressed | 2.20 |

#### Table S1. Continued

| | | | | |
|---|---|---|---|---|
| gene:Os08g0206600 | Os08g0206600 | LOC_Os08g10570.1 | bifunctional purine biosynthesis protein purH, putative, expressed | 2.19 |
| OsMT1g | Os12g0571000 | LOC_Os12g38290.1 | metallothionein, putative, expressed | 2.19 |
| Stt3a | Os04g0675500 | LOC_Os04g57890.1 | oligosaccharyl transferase, putative, expressed | 2.18 |
| gene:Os06g0634000 | Os06g0634000 | LOC_Os06g42754.1 | expressed protein | 2.18 |
| HWH1 | Os02g0621800 | LOC_Os02g40840.1 | alcohol oxidase, putative, expressed | 2.18 |
| gene:Os02g0206700 | Os02g0206700 | LOC_Os02g11640.1 | UDP-glucoronosyl and UDP-glucosyl transferase, putative, expressed | 2.15 |
| gene:Os10g0189600 | Os10g0189600 | LOC_Os10g11200.1 | aminotransferase, classes I and II, domain containing protein, expressed | 2.15 |
| gene:Os07g0563000 | Os07g0563000 | LOC_Os07g37580.1 | diacylglycerol kinase, putative, expressed | 2.14 |
| CAL1 | Os02g0629800 | LOC_Os02g41904.1 | DEF7 - Defensin and Defensin-like DEFL family, expressed | 2.14 |
| gene:Os05g0111200 | Os05g0111200 | LOC_Os05g02060.1 | mitochondrial import inner membrane translocase subunit Tim17, putative, expressed | 2.12 |
| gene:Os06g0652600 | Os06g0652600 | LOC_Os06g44280.1 | retrotransposon protein, putative, Ty3-gypsy subclass, expressed | 2.12 |
| gene:Os02g0705400 | Os02g0705400 | LOC_Os02g47650.1 | universal stress protein domain containing protein, putative, expressed | 2.12 |
| OsEXPA11 | Os01g0274500 | LOC_Os01g16770.1 | expansin precursor, putative, expressed | 2.11 |
| RBBI3-3 | Os01g0124401 | LOC_Os01g03360.1 | BBTI5 - Bowman-Birk type bran trypsin inhibitor precursor, expressed | 2.11 |
| gene:Os06g0716100 | Os06g0716100 | LOC_Os06g50230.1 | expressed protein | 2.10 |
| gene:Os01g0300200 | Os01g0300200 | LOC_Os01g19450.1 | ATP-citrate synthase subunit 1, putative, expressed | 2.09 |
| NAC131 | Os12g0123700 | LOC_Os12g03040.1 | no apical meristem protein, putative, expressed | 2.09 |
| gene:Os03g0434400 | Os03g0434400 | LOC_Os03g32040.1 | phenazine biosynthesis protein, putative, expressed | 2.09 |
| GRF1 | Os02g0776900 | LOC_Os02g53690.1 | growth regulating factor protein, putative, expressed | 2.09 |
| D11 | Os04g0469800 | LOC_Os04g39430.1 | cytochrome P450, putative, expressed | 2.09 |
| OsLTP1.9 | Os11g0115100 | LOC_Os11g02350.1 | LTPL25 - Protease inhibitor/seed storage/LTP family protein precursor, expressed | 2.09 |
| OsPTR3_1 | Os10g0470700 | LOC_Os10g33210.1 | peptide transporter PTR3-A, putative, expressed | 2.08 |
| gene:Os05g0318600 | Os05g0318600 | LOC_Os05g25430.1 | receptor-like protein kinase At3g46290 precursor, putative, expressed | 2.08 |
| AP2/EREBP129 | Os01g0141000 | LOC_Os01g04800.1 | B3 DNA binding domain containing protein, expressed | 2.08 |
| gene:Os01g0947000 | Os01g0947000 | LOC_Os01g71860.1 | glycosyl hydrolases family 17, putative, expressed | 2.08 |
| gene:Os03g0129800 | Os03g0129800 | LOC_Os03g03730.1 | regulatory protein, putative, expressed | 2.07 |
| gene:Os11g0199801 | Os11g0199801 | LOC_Os11g09350.1 | expressed protein | 2.06 |
| gene:Os10g0524700 | Os10g0524700 | LOC_Os10g38090.1 | cytochrome P450, putative, expressed | 2.06 |
| gene:Os01g0693300 | Os01g0693300 | LOC_Os01g49820.1 | lipid phosphatase protein, putative, expressed | 2.06 |
| RAC1 | Os01g0229400 | LOC_Os01g12900.1 | ras-related protein, putative, expressed | 2.04 |
| OsUGT98B1 | Os01g0176000 | LOC_Os01g08090.1 | flavonol-3-O-glycoside-7-O-glucosyltransferase 1, putative, expressed | 2.03 |
| Osr9-LOX1 | Os03g0699700 | LOC_Os03g49260.1 | lipoxygenase, putative, expressed | 2.03 |
| OsTHI29 | Os03g0247200 | LOC_Os03g14300.1 | THION29 - Plant thionin family protein precursor, expressed | 2.01 |
| gene:Os06g0290701 | Os06g0290701 | None | | 2.01 |
| gene:Os06g0676700 | Os06g0676700 | LOC_Os06g46340.1 | glycosyl hydrolase, family 31, putative, expressed | 2.01 |

### Table S2. Down-regulated genes of 10 μM TIS108-treated rice

| Gene symbol | RAP ID | MSU ID | Description | Fold change |
|---|---|---|---|---|
| gene:Os05g0122600 | Os05g0122600 | LOC_Os05g03120.1 | retrotransposon protein, putative, unclassified, expressed | -723.50 |
| EIP11 | Os08g0476300 | LOC_Os08g37130.1 | oxidoreductase, short chain dehydrogenase/reductase family domain containing protein, expressed | -473.03 |
| gene:Os04g0488600 | Os04g0488600 | LOC_Os04g41150.1 | DUF565 domain containing protein, putative, expressed | -459.00 |
| gene:Os05g0393400 | Os05g0393400 | LOC_Os05g32680.1 | PAC, putative, expressed | -194.64 |
| gene:Os03g0335500 | Os03g0335500 | LOC_Os03g21730.1 | receptor-like protein kinase precursor, putative, expressed | -147.11 |
| gene:Os07g0178950 | Os07g0178950 | None | | -57.10 |
| gene:Os10g0487400 | Os10g0487400 | LOC_Os10g34590.1 | zinc finger, C3HC4 type domain containing protein, expressed | -37.62 |
| gene:Os04g0632400 | Os04g0632400 | None | #N/A | -26.10 |
| prx7 | Os07g0157000 | LOC_Os07g06300.1 | ethylene-insensitive protein 2, putative, expressed | -24.13 |
| RR7 | Os07g0449700 | LOC_Os07g26720.1 | OsRR7 type-A response regulator, expressed | -22.52 |
| gene:Os07g0156467 | Os07g0156467 | LOC_Os07g06190.1 | ethylene-insensitive protein 2, putative, expressed | -21.16 |
| gene:Os07g0633100 | Os07g0633100 | LOC_Os07g43940.1 | X8 domain containing protein, expressed | -18.72 |
| OMT7 | Os03g0708100 | LOC_Os03g50040.1 | phytanoyl-CoA dioxygenase, putative, expressed | -16.91 |
| gene:Os11g0495950 | Os11g0495950 | LOC_Os11g30310.1 | reticuline oxidase-like protein precursor, putative, expressed | -15.77 |
| OsPKS16 | Os10g0158400 | LOC_Os10g07040.1 | chalcone synthase, putative, expressed | -14.00 |
| gene:Os07g0107583 | Os07g0107583 | None | | -13.57 |
| gene:Os05g0435300 | Os05g0435300 | LOC_Os05g35960.1 | expressed protein | -13.55 |
| gene:Os12g0173125 | Os12g0173125 | None | | -12.03 |
| gene:Os02g0593700 | Os02g0593700 | LOC_Os02g38050.1 | joka2, putative, expressed | -11.51 |
| CHS_2 | Os05g0212900 | LOC_Os05g12210.1 | chalcone synthase, putative, expressed | -10.76 |
| gene:Os12g0614201 | Os12g0614201 | None | | -10.19 |
| OsPKS07 | Os05g0213100 | LOC_Os05g12240.1 | chalcone synthase, putative, expressed | -8.57 |
| gene:Os01g0164075 | Os01g0164075 | LOC_Os01g07030.1 | POEI40 - Pollen Ole e I allergen and extensin family protein precursor, expressed | -8.39 |
| OsbHLH130 | Os12g0589000 | LOC_Os12g39850.1 | helix-loop-helix DNA-binding domain containing protein, expressed | -7.82 |
| OsPME9 | Os02g0783000 | LOC_Os02g54190.1 | pectinesterase, putative, expressed | -7.50 |
| PRX118 | Os08g0302000 | LOC_Os08g20730.1 | peroxidase precursor, putative, expressed | -7.50 |
| OSH1 | Os03g0727000 | LOC_Os03g51690.1 | Homeobox domain containing protein, expressed | -6.45 |
| gene:Os11g0156000 | Os11g0156000 | LOC_Os11g05740.1 | B3 DNA binding domain containing protein, expressed | -6.33 |
| OsCCR_1 | Os08g0277200 | LOC_Os08g17500.1 | cinnamoyl-CoA reductase, putative, expressed | -6.10 |
| OsalphaCA5 | Os08g0424100 | LOC_Os08g32840.1 | bifunctional monodehydroascorbate reductase and carbonic anhydrasenectarin-3 precursor, putative, e | -5.68 |
| gene:Os11g0156000 | Os11g0156000 | LOC_Os11g05740.1 | B3 DNA binding domain containing protein, expressed | -5.33 |
| GLP8-4 | Os08g0189300 | LOC_Os08g08980.1 | cupin domain containing protein, expressed | -5.28 |
| gene:Os12g0614100 | Os12g0614100 | LOC_Os12g41970.1 | lipase class 3 family protein, putative, expressed | -5.05 |
| OsSub33 | Os04g0120300 | LOC_Os04g02980.1 | OsSub33 - Putative Subtilisin homologue, expressed | -4.88 |
| OsLLA8 | Os10g0191100 | LOC_Os10g11370.1 | LTPL87 - Protease inhibitor/seed storage/LTP family protein precursor, putative, expressed | -4.65 |
| gene:Os11g0289700 | Os11g0289700 | LOC_Os11g18570.1 | cytochrome P450, putative, expressed | -4.63 |
| gene:Os01g0163450 | Os01g0163450 | None | | -4.28 |
| GLP8-3 | Os08g0189200 | LOC_Os08g08970.1 | Cupin domain containing protein, expressed | -3.90 |
| OsCrRLK1L1 | Os03g0759600 | LOC_Os03g55210.1 | TKL_IRAK_CrRLK1L-1.1 - The CrRLK1L-1 subfamily has homology to the CrRLK1L homolog, expressed | -3.88 |
| gene:Os02g0124599 | Os02g0124599 | LOC_Os02g03210.1 | FAD-binding and arabino-lactone oxidase domains containing protein, putative, expressed | -3.75 |
| gene:Os10g0538450 | Os10g0538450 | None | | -3.73 |
| gene:Os06g0169001 | Os06g0169001 | LOC_Os06g07250.1 | jacalin-like lectin domain containing protein, expressed | -3.69 |
| gene:Os02g0184800 | Os02g0184800 | None | | -3.50 |
| OsTIP4_1 | Os01g0232000 | LOC_Os01g13120.1 | aquaporin protein, putative, expressed | -3.44 |
| OsERF#039 | Os01g0200600 | LOC_Os01g10370.1 | AP2 domain containing protein, expressed | -3.35 |
| gene:Os10g0450800 | Os10g0450800 | LOC_Os10g31320.1 | retrotransposon protein, putative, unclassified, expressed | -3.30 |
| OsIAA9 | Os02g0805100 | LOC_Os02g56120.1 | OsIAA9 - Auxin-responsive Aux/IAA gene family member, expressed | -3.25 |
| gene:Os09g0364800 | Os09g0364800 | LOC_Os09g20000.1 | heavy metal-associated domain containing protein, expressed | -3.24 |
| OsPP2C49 | Os05g0457200 | LOC_Os05g38290.1 | protein phosphatase 2C, putative, expressed | -3.24 |
| gene:Os05g0148800 | Os05g0148800 | LOC_Os05g05610.1 | expressed protein | -3.24 |
| gene:Os11g0303300 | Os11g0303300 | LOC_Os11g19780.1 | O-methyltransferase ZRP4, putative | -3.18 |
| gene:Os01g0191200 | Os01g0191200 | LOC_Os01g09540.1 | HAD superfamily phosphatase, putative, expressed | -3.15 |
| gene:Os03g0281466 | Os03g0281466 | LOC_Os03g17310.1 | calcium-transporting ATPase, endoplasmic reticulum-type, putative, expressed | -3.10 |
| gene:Os06g0179500 | Os06g0179500 | LOC_Os06g08120.1 | plant protein of unknown function domain containing protein, expressed | -3.10 |
| gene:Os01g0228450 | Os01g0228450 | None | | -3.10 |
| gene:Os05g0588900 | Os05g0588900 | LOC_Os05g51130.1 | mitochondrial chaperone BCS1, putative, expressed | -3.05 |
| gene:Os02g0596300 | Os02g0596300 | LOC_Os02g38290.1 | cytochrome P450, putative, expressed | -2.98 |
| gene:Os11g0635300 | Os11g0635300 | LOC_Os11g41680.1 | cytochrome P450, putative, expressed | -2.93 |
| OsSAUR33 | Os08g0452500 | LOC_Os08g35110.1 | OsSAUR33 - Auxin-responsive SAUR gene family member, expressed | -2.91 |
| gene:Os03g0740200 | Os03g0740200 | LOC_Os03g52940.1 | expressed protein | -2.87 |
| gene:Os04g0644100 | Os04g0644100 | LOC_Os04g55120.1 | jp18, putative, expressed | -2.84 |
| gene:Os09g0412700 | Os09g0412700 | LOC_Os09g24620.1 | expressed protein | -2.81 |
| gene:Os02g0542600 | Os02g0542600 | None | | -2.74 |
| OsTPP1 | Os02g0661100 | LOC_Os02g44230.1 | CPuORF22 - conserved peptide uORF-containing transcript, expressed | -2.71 |
| gene:Os03g0700450 | Os03g0700450 | None | | -2.70 |
| gene:Os06g0567433 | Os06g0567433 | None | | -2.64 |
| OsDXS | Os07g0190000 | LOC_Os07g09190.1 | transketolase, putative, expressed | -2.61 |
| gene:Os04g0400600 | Os04g0400600 | LOC_Os04g32820.1 | expressed protein | -2.61 |

#### Table S2. Continued

| | | | | |
|---|---|---|---|---|
| OsTPP1 | Os02g0661100 | LOC_Os02g44230.1 | CPuORF22 - conserved peptide uORF-containing transcript, expressed | -2.60 |
| CHS_1 | Os04g0103900 | LOC_Os04g01354.1 | chalcone synthase, putative, expressed | -2.58 |
| gene:Os12g0614200 | Os12g0614200 | LOC_Os12g41980.1 | lipase class 3 family protein, putative, expressed | -2.54 |
| OsOPR9 | Os01g0370000 | LOC_Os01g27240.1 | 12-oxophytodienoate reductase, putative, expressed | -2.53 |
| gene:Os04g0538400 | Os04g0538400 | LOC_Os04g45520.1 | integral membrane protein, putative, expressed | -2.49 |
| gene:Os01g0588100 | Os01g0588100 | LOC_Os01g40560.1 | hypersensitive-induced response protein, putative, expressed | -2.41 |
| gene:Os01g0916100 | Os01g0916100 | LOC_Os01g68740.1 | keratin, type I cytoskeletal 9, putative, expressed | -2.40 |
| gene:Os01g0926400 | Os01g0926400 | LOC_Os01g70180.1 | exostosin family domain containing protein, expressed | -2.39 |
| OsPUB39 | Os06g0248500 | LOC_Os06g13870.1 | U-box protein CMPG1, putative, expressed | -2.38 |
| OsXTH13 | Os02g0280200 | LOC_Os02g17880.1 | glycosyl hydrolases family 16, putative, expressed | -2.37 |
| gene:Os09g0248900 | Os09g0248900 | LOC_Os09g07440.1 | retrotransposon protein, putative, unclassified, expressed | -2.35 |
| gene:Os03g0242300 | Os03g0242300 | LOC_Os03g13870.1 | expressed protein | -2.34 |
| gene:Os07g0638801 | Os07g0638801 | None | | -2.33 |
| ICS1 | Os09g0361500 | LOC_Os09g19734.1 | isochorismate synthase 1, chloroplast precursor, putative, expressed | -2.28 |
| gene:Os02g0628200 | Os02g0628200 | LOC_Os02g41780.1 | transporter-related, putative, expressed | -2.23 |
| gene:Os04g0630400 | Os04g0630400 | LOC_Os04g53810.1 | leucoanthocyanidin reductase, putative, expressed | -2.23 |
| Os-PHT2 | Os06g0185300 | LOC_Os06g08610.1 | transferase family protein, putative, expressed | -2.19 |
| gene:Os04g0578300 | Os04g0578300 | LOC_Os04g48870.1 | nitrilase-associated protein, putative, expressed | -2.19 |
| prx105 | Os07g0638600 | LOC_Os07g44460.1 | peroxidase precursor, putative, expressed | -2.19 |
| OsMC8 | Os03g0389100 | LOC_Os03g27190.1 | ICE-like protease p20 domain containing protein, putative, expressed | -2.13 |
| gene:Os04g0550866 | Os04g0550866 | None | | -2.12 |
| gene:Os01g0369950 | Os01g0369950 | None | | -2.11 |
| OsCDAP2 | Os07g0162700 | LOC_Os07g06860.1 | gibberellin receptor GID1L2, putative, expressed | -2.11 |
| OsPIP2_4 | Os04g0521100 | LOC_Os04g44060.1 | aquaporin protein, putative, expressed | -2.10 |
| gene:Os01g0700500 | Os01g0700500 | LOC_Os01g50490.1 | cytochrome P450, putative, expressed | -2.09 |
| gene:Os01g0164300 | Os01g0164300 | LOC_Os01g07060.1 | POEI43 - Pollen Ole e I allergen and extensin family protein precursor, putative, expressed | -2.07 |
| gene:Os03g0760500 | Os03g0760500 | LOC_Os03g55260.1 | cytochrome P450, putative, expressed | -2.06 |
| prx68 | Os05g0135000 | LOC_Os05g04450.1 | peroxidase precursor, putative, expressed | -2.06 |
| gene:Os03g0134500 | Os03g0134500 | LOC_Os03g04190.1 | cytochrome P450, putative, expressed | -2.06 |
| gene:Os06g0199700 | Os06g0199700 | LOC_Os06g09920.1 | expressed protein | -2.05 |
| gene:Os10g0162842 | Os10g0162842 | None | | -2.05 |
| gene:Os05g0355700 | Os05g0355700 | LOC_Os05g28770.1 | GCRP9 - Glycine and cysteine rich family protein precursor, expressed | -2.04 |
| gene:Os06g0133600 | Os06g0133600 | LOC_Os06g04250.1 | phosphate-induced protein 1 conserved region domain containing protein, expressed | -2.04 |
| gene:Os12g0135800 | Os12g0135800 | LOC_Os12g04150.1 | alpha/beta hydrolase fold, putative, expressed | -2.04 |
| OsIAA20 | Os06g0166500 | LOC_Os06g07040.1 | OsIAA20 - Auxin-responsive Aux/IAA gene family member, expressed | -2.03 |
| NaT | Os05g0382200 | LOC_Os05g31730.1 | transporter, monovalent cation:proton antiporter-2 family, putative, expressed | -2.02 |
| gene:Os11g0644700 | Os11g0644700 | LOC_Os11g42500.1 | dirigent, putative, expressed | -2.01 |
| OsStr7 | Os12g0428000 | LOC_Os12g24020.1 | rhodanese-like domain containing protein, putative, expressed | -2.01 |

#### Table S3. Expression of strigolactone- and tillering-related genes

| Genes | RAP ID | Variant | Fold change | FDR p-value |
|---|---|---|---|---|
| D27 9-cis/all-trans-P-carotene isomerase | Os11g0587000 | D27_1 | -1.82 | 0.69 |
| | | D27_2 | -1.12 | 1.00 |
| D10 Carotenoid cleavage dioxygenase8 | Os01g0746400 | D10_1 | -1.68 | 0.07 |
| D17 Carotenoid cleavage dioxygenase7 | Os04g0550600 | HED1_1 | -1.75 | 0.05 |
| MAX1 (Os5100) Cytochrome P450 | Os06g0565100 | Os5100_1 | -1.31 | 0.61 |
| MAX1 (Os1900) Cytochrome P450 | Os02g0221900 | Os1900_1 | -66.14 | 0.45 |
| | | Os1900_2 | -1.38 | 1.00 |
| | | Os1900_3 | 1.06 | 1.00 |
| MAX1 (Os900) Cytochrome P450 | Os01g0700900 | SLB1_1 | -1.07 | 1.00 |
| | | SLB1_2 | -1.19 | 1.00 |
| MAX1 (Os1400) Cytochrome P450 | Os01g0701400 | SLB2_1 | -2.08 | 0.69 |
| D3 F-box | Os06g0154200 | D3_1 | 1.33 | 0.67 |
| | | D3_2 | 1.33 | 0.53 |
| D14 α / β -Hydrolase | Os03g0203200 | D14_1 | 1.39 | 1.00 |
| | | D14_2 | -1.02 | 1.00 |
| D53 Class I Clp ATPase | Os11g0104300 | D53_1 | -1.39 | 0.24 |
| | | D53_2 | -1.10 | 1.00 |
| FC1 | Os03g0706500 | FC1_1_1 | -3.19 | 1.00 |
| REP1 | Os09g0410500 | REP1_1 | -1.03 | 1.00 |
| OsIPA1 | Os08g0509600 | WFP_1 | 1.18 | 1.00 |
| NSP1 | Os03g0408600 | NSP1_1 | -1.21 | 0.83 |
| NSP2 | Os03g0263300 | NSP2_1 | -1.19 | 0.97 |

Table S4: Primer lists for RT-qPCR.

| Primer name | Primer sequence | Organism |
|---|---|---|
| UbiQ | F:GCCCAAGAAGAAGATCAAGAAC<br>R: AGATAACAACGGAAGCATAAAAGTC | Rice Nipponbare |
| OsMAX1-900 | F:ATTGTCAGCGATCCACTTC<br>R:GCGCCGTTCTTGAAATTG | Rice Nipponbare |
| OsMAX1-1400 | F:GGCAGGTGCTCAAGAGGATT<br>R:TTTTGTCCATCTGTCCCCCG | Rice Nipponbare |
| OsMAX1-1900 | F:GTTCCCCATAGGCCACCTTC<br>R:GCATTGGCCACAATCACCAG | Rice Nipponbare |
| OsMAX1-5100 | F:GTGATAAAGGAGGCGATGAG<br>R:CTTTGGGAGTGTGTAGCC | Rice Nipponbare |
| OsRubQ1 | F:GGGTTCACAAGTCTGCCTATTTG<br>R: ACGGGACACGACCAAGGA | AMF – *R. irregularis* |
| OsPT11 | F: GAGAAGTTCCCTGCTTCAAGCA<br>R: CATATCCCAGATGAGCGTATCATG | AMF – *R. irregularis* |
